# Spiking recurrent neural networks represent task-relevant neural sequences in rule-dependent computation

**DOI:** 10.1101/2021.01.21.427464

**Authors:** Xiaohe Xue, Michael M. Halassa, Zhe S. Chen

**Affiliations:** Courant Institute of Mathematical Sciences, New York University, New York, New York, United States of America; Department of Brain and Cognitive Sciences, Massachusetts Institute of Technology, Cambridge, Massachusetts, United States of America; Department of Psychiatry, New York University School of Medicine, New York, New York, United States of America; Department of Neuroscience and Physiology, New York University School of Medicine, New York, New York, United States of America; Neuroscience Institute, New York University School of Medicine, New York, New York, United States of America

## Abstract

Prefrontal cortical neurons play in important roles in performing rule-dependent tasks and working memory-based decision making. Motivated by experimental data, we develop an excitatory-inhibitory spiking recurrent neural network (SRNN) to perform a rule-dependent two-alternative forced choice (2AFC) task. We imposed several important biological constraints onto the SRNN, and adapted the spike frequency adaptation (SFA) and SuperSpike gradient methods to update the network parameters. These proposed strategies enabled us to train the SRNN efficiently and overcome the vanishing gradient problem during error back propagation through time. The trained SRNN produced rule-specific tuning in single-unit representations, showing rule-dependent population dynamics that strongly resemble experimentally observed data in rodent and monkey. Under varying test conditions, we further manipulated the parameters or configuration in computer simulation setups and investigated the impacts of rule-coding error, delay duration, weight connectivity and sparsity, and excitation/inhibition (E/I) balance on both task performance and neural representations. Overall, our modeling study provides a computational framework to understand neuronal representations at a fine timescale during working memory and cognitive control.

**Author Summary:** Working memory and decision making are fundamental cognitive functions of the brain, but the circuit mechanisms of these brain functions remain incompletely understood. Neuroscientists have trained animals (rodents or monkeys) to perform various cognitive tasks while simultaneously recording the neural activity from specific neural circuits. To complement the experimental investigations, computational modeling may provide an alternative way to examine the neural representations of neuronal assemblies during task behaviors. Here we develop and train a spiking recurrent neural network (SRNN) consisting of balanced excitatory and inhibitory neurons to perform the rule-dependent working memory tasks Our computer simulations produce qualitatively similar results as the experimental findings. Moreover, the imposed biological constraints on the trained network provide additional channel to investigate cell type-specific population responses, cortical connectivity and robustness. Our work provides a computational platform to investigate neural representations and dynamics of cortical circuits a fine timescale during complex cognitive tasks.

## Introduction

A biological brain is large-scale neuronal network with recurrent connections that performs computations in complex task behaviors. In recent years, recurrent neural networks (RNNs) have been widely used for modeling a wide range of neural circuits, such as the prefrontal cortex (PFC), parietal cortex, and primary motor cortex (M1), in various cognitive and motor tasks [1-9]. Varying types of assumptions have been made in different models: leaky integrate-and-fire (LIF) vs. conductance-based compartment model, rate vs. spiking-based model. Spikes are the fundamental language that brain uses to represent and communicate information. Spiking neural networks (SNNs) are inspired by information processing in the brain, and attempt to mimic the mechanistic operation and computation among spiking neurons [10]. Although SNNs are derived from biologically inspired neurons, neural plasticity or learning for SNNs have been less efficient (i.e., slow convergence) compared to the rate-based models [11-14]. In recent years, many methods have been developed for improving the learning speed of feedforward and recurrent SNNs [15-18].

Working memory and attention control are two fundamental functions in cognition and decision making. The two-alternative forced choice (2AFC) task is a classic experiment in psychophysics in electrophysiology to measure the working memory and cognitive control capacity. In light of recent experiments [19,20], PFC neurons are found to show rule-specific and timing-specific neuronal responses during the delay (i.e., working memory) period, emerging rule-specific neural sequences (Fig. 1). Sequentially activated neuronal activities (“neural sequences”) have been widely observed in neural assemblies in cortical areas during working memory or decision making [19,21,22]. In the literature, neural sequences have been generated by RNNs [4, 23]. However, these sequences were trained by supervised learning; it remains unknown whether the rule-specific neural representations or sequences can emerge from a trained RNN. It is also unclear how these neural representations change as a function of network excitation and inhibition. Inspired by these questions, we developed an excitatory-inhibitory SRNN model that satisfies Dale’s principle [7, 24], and we adapted a SuperSpike surrogate gradient learning algorithm [17] and SFA method [25, 26] to train the SRNN. Unlike the rate-based RNN, SRNN can provide millisecond resolution to characterize neuronal tuning.

**Fig 1.**
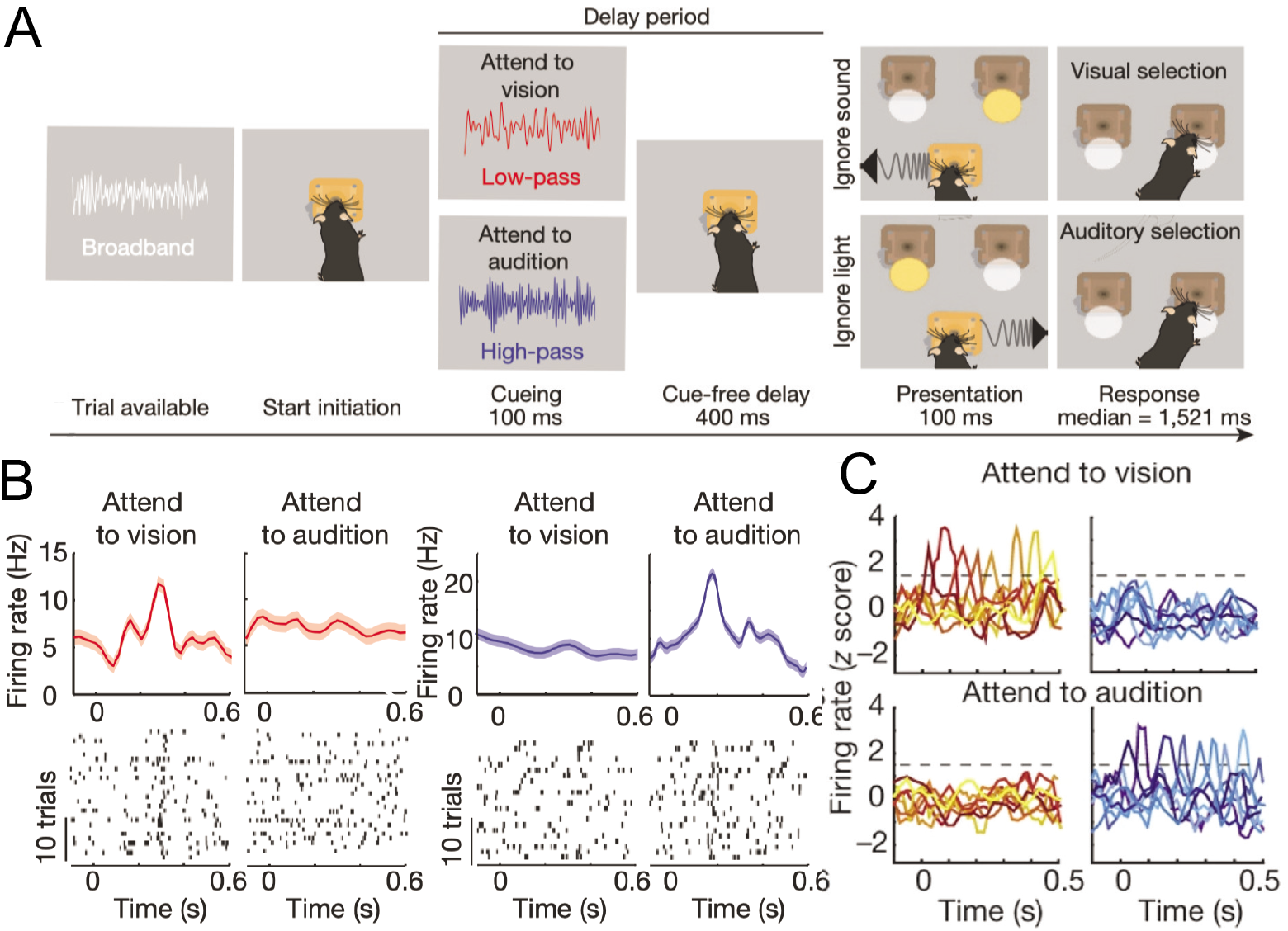
Rule-specific sequential PFC activity emerges during a rodent working memory task. (A) Schematic of 2AFC task. (B) Representative peri-stimulus time histogram (PSTH) and rasters for mouse PFC neurons with specific tuning of attending to vision (rule 1) or attending to audition (rule 2). (C) Examples of tuning peaks from mouse PFC neurons across multiple sessions (Figures reproduced from [19] with permission, © Springer Nature, 2017).

The contributions of this paper are threefold. First, to alleviate the vanishing gradient problem, we adapted the SuperSpike surrogate gradient algorithm to learn excitatory-inhibitory SRNN parameters. These combined computational strategy is key to efficiently train the SRNN for a delayed working memory task. Second, we showed the trained SRNN produced emergent task-relevant neural representations of excitatory and inhibitory neurons, such as neural sequences and oscillatory spiking during the delay period. Our computer model prediction match the published experimental data. Third, we manipulated and quantified the impact of the excitatory-to-inhibitory (E/I) balance, delay duration, and cortical connectivity on task performance and neural representations. Overall, our large-scale SRNN modeling and simulations provide a computational framework to study task representations and dynamics in a wide range of cognitive tasks.

## Methods

### Rule-dependent working memory

In the 2AFC task, animals were trained to initiate a trial upon receiving broadband white noise. Upon successful initiation, the white noise was immediately replaced by either low-pass or high-pass filtered noise for 100 ms to indicate the rule (i.e., attending to vision vs. audition). This is was followed by a delay period lasting ~ 400 ms before the target stimuli presentation. Animals received a reward if their behavior led to a correct rule-specific response at two specific response ports (Fig. 1A, [19]). In an extension of 2AFC task, animals were trained to respond to four response ports instead of two response ports (known as 4AFC task), whereas the other task phases remained unchanged. Responses were scored as correct or one of three different error types (executive error, sensory error, or both). All animal experiments were performed according to the guidelines of the US National Institutes of Health and the Institutional Animal Care and Use Committee at the New York University Langone Medical Center.

### Computer simulation paradigm based on SRNN

To mimic the rule-dependent computation in the PFC circuit, we set up a computational model for performing the cognitive task. The complete model consisted of three independent components: an input encoder, a SRNN and an output decoder, as shown in Fig. 2A. The role of input encoder was to transform the analog input signal into spikes. We used five leaky LIF neurons for representing the 2AFC task: one neuron for generating cueing or rule inputs, and the other four neurons for encoding distinct sensory cues (e.g., Vision/Left, Vision/Right, Audition/Left, Audition/Right as in [19], Fig. 2B). Therefore, two rules in this task were to attend to vision (Rule 1) versus attend to audition (rule 2), where the context cues were represented by different cues (cue 1 versus cue 2). The role of output decoder was to integrate and transform the spikes from the SRNN into an analog signal for final (binary) decision. We used two LIF neurons for the 2AFC task. The final voltage values of readout neurons were interpreted as the probabilities of binary decision.

**Fig 2.**
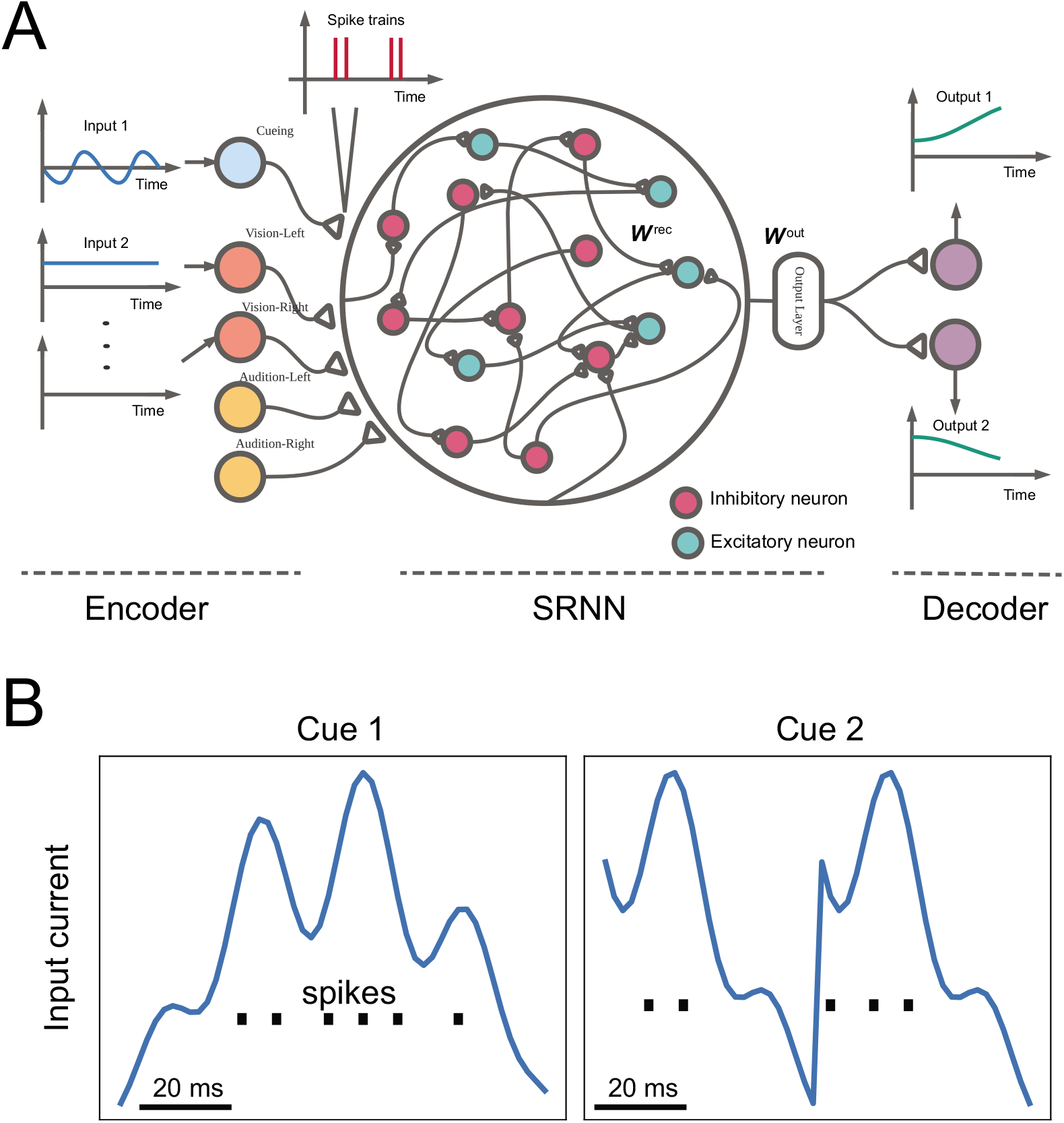
Schematic of computer simulations: an input encoder, a SRNN and an output decoder. (A) The SRNN consisting of both excitatory and inhibitory spiking neurons, receives the input spike trains converted from an encoder, and generates the output voltage and spike trains that are fed into a decoder, which generates the final decision. The neurons are fully connected with recurrent weights ***W***^rec^, and the readout layer was parameterized by ***W***^out^. (B) Illustration of encoding two cueing current inputs into spikes.

To mimic the behavioral task in rodent experiments, we used the following computer simulation setup:

- 200-ms fixation period, during which only white noise was supplied.
- 100-ms cueing period, during which the spike trains were generated by the cueing encoder and sent to the SRNN.
- 400-ms cueing delay period, during which only white noise was supplied.
- 100-ms stimulus presentation period, during which two spike trains were represented: one from the visual channel, and the other from the auditory channel.
- Response period, during which the cost function was computed at the end of the presentation period, and the network back-propagated the error and computed the gradient.

For each rule condition, we simulated independent Monte Carlo trials based on different initial conditions. In what follows, we present the mathematical details and implementation of each component. The mathematical symbols and notations are summarized in Table 1.

**Table 1.**
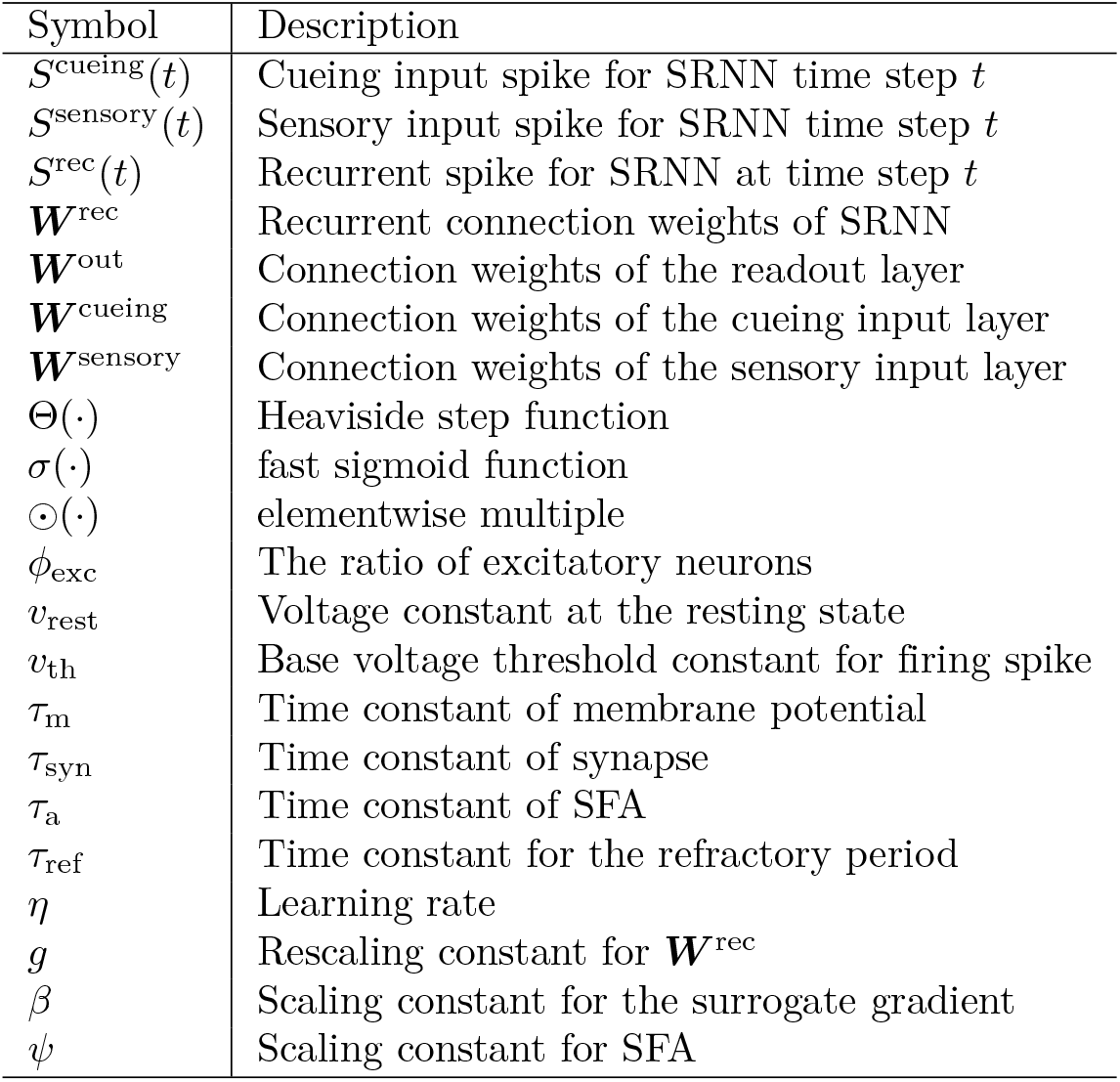
A summary of symbols and notations in SRNN.

#### Mathematical description of SRNN

The structure of SRNN had a similar form as the rate-based RNNs [7], except that each unit of the SRNN consisted of a biologically-constrained LIF neuron. All neurons were connected through a recurrent weight matrix ***W***^rec^, and the SRNN received cueing/sensory inputs from the encoders via ***W***^cueing^ and ***W***^sensory^, and produced outputs through the output weight matrix. The basic dynamics of neurons in SRNN was described as follows:

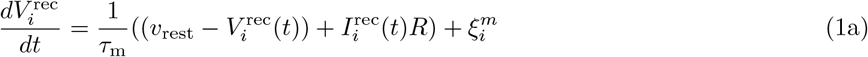

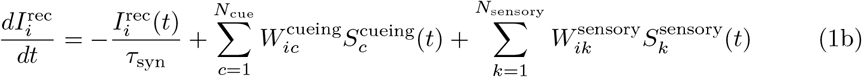

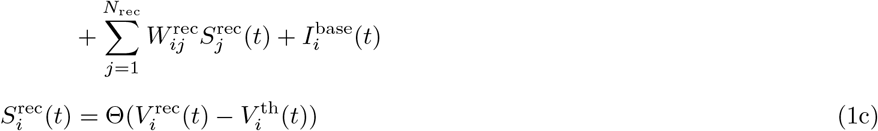

Each neuron was modeled by a leaky integrator, where Θ denote the Heaviside step function. In Eq. 1a, the dynamics of membrane potential 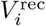 is driven by synaptic current *I_i_* and zero-mean white noise 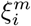, and compared with its resting state *υ*_rest_. In Eq. 1b, the synaptic current current *I_i_* was modeled by a linear sum of all presynaptic spikes through three associated weight matrices 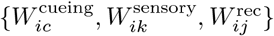. Spike trains 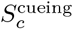 and 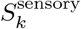 originated from the *c*-th cueing encoder and the *K*-the sensory encoder, respectively; and *S*^rec^ represented the spike trains from other recurrently connected neurons. The current 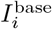 was injected to maintain a baseline firing rate. Finally, a spike 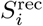 would fire when *V_i_* was above the spike threshold 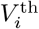. Right after firing, 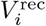 was reset to membrane potential resting state *v*_rest_ with a short refractory period *τ*_ref_.

Additionally, we added random noise to the SRNN to emulate the noisy nervous system. Two sources of noise were considered such that the SRNN still maintained a reasonable level of neuronal firing rate even in the absence of external inputs. The first one was the voltage noise ξ^*m*^ (Eq. 1a), which was sampled from a zero-mean normal distribution. The second source of noise was the baseline current noise 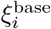 (Eq. 2), which was also sampled from a zero-mean normal distribution at every step by following a random-walk model for the baseline current 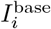:

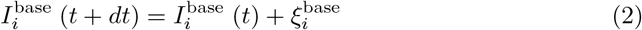

Our SRNN has several important features that will be discussed later: (i) the SRNN consisted of excitatory and inhibitory neurons that follow Dale’s principle; (ii) the SRNN implemented an efficient SuperSpike algorithm [17] to alleviate the vanishing gradient problem; (iii) the SRNN employed a SFA strategy to strengthen the memory capacity during the delay period [25]; and (iv) the SRNN used a regularized cost function that penalized the excessive spikes in the neuronal population (thereby imposing a sparsity constraint).

#### Network initialization

We initialized the neuronal states and neuronal baseline firing rates using the following procedure. First, we uniformly randomized the initial membrane potential for every neurons within a reasonable range (between the membrane potential resting state *υ*_rest_ and the baseline spiking threshold *υ*_th_) at each trial. This range was chosen to improve the robustness of the model. Second, the initial baseline current for each neuron was sampled from a truncated normal distribution, which did not varied across different trials (Table 3).

**Table 2.** Parameters used in the encoder and decoder.

| Symbol | Description |
| --- | --- |
| $V^{\text{encoder}}(t=0)$ | 0.0 mV |
| $V^{\text{decoder}}(t=0)$ | 0.0 mV |
| $v_{\text{rest}}^{\text{encoder}}$ | 0.0 mV |
| $v_{\text{rest}}^{\text{decoder}}$ | 0.0 mV |
| $v_{\text{th}}^{\text{encoder}}$ | 1.0 mV |
| $v_{\text{th}}^{\text{decoder}}$ | 1.0 mV |
| $I^{\text{encoder}}(t=0)$ | 0.0 |
| $I^{\text{decoder}}(t=0)$ | 0.0 |
| $R^{\text{encoder}}$ | 1 |
| $R^{\text{decoder}}$ | 1 |
| $\tau_{\text{m}}^{\text{encoder}}$ | 5 ms |
| $\tau_{\text{m}}^{\text{decoder}}$ | 5 ms |
| $\tau_{\text{syn}}^{\text{decoder}}$ | 20 ms |

**Table 3.** Hyperparameters used in the SRNN.

| Symbol | Description |
| --- | --- |
| $I^{\text{base}}(t=0)$ | $0.041 \times \mathcal{N}(\mu=2, \sigma^2=0.01)$, truncated in $[1.8, 2.4]$ |
| $\xi^{\text{base}}(t)$ | $\mathcal{N}(\mu=0, \sigma^2=4 \times 10^{-6})$ |
| $\xi^m(t)$ | $\mathcal{N}(\mu=0, \sigma^2=16)$ |
| $v_{\text{rest}}$ | -65 mV |
| $v_{\text{th}}$ | -50 mV |
| $\phi_{\text{exc}}$ | 80% |
| $dt$ | 2 ms |
| $\beta$ | 1000 |
| $R$ | $10^{-2}$ |
| $\tau_{\text{mem}}$ | 20 ms |
| $\tau_{\text{syn}}$ | 35 ms for excitatory neurons, 40 ms for inhibitory neurons |
| $\tau_{\text{a}}$ | 400 ms |
| $\tau_{\text{ref}}$ | 6 ms |
| $\eta$ | $3 \times 10^{-4}$ |
| $g$ | 1.5 |
| $\psi$ | 1.6 |
| $\lambda$ | $10^{-9}$ |

All the connection weights of the SRNN were firstly initialized via the Glorot method [27]. Given a matrix with the size of fan_in_ × fan_out_, every connection weight *w*_init_ was sampled from a uniform distribution as follows

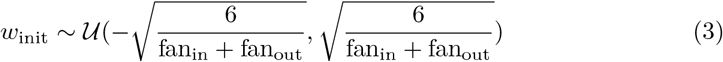

#### Encoder and decoder

Both the encoder and decoder (Fig. 2A) followed the same LIF neuron structure; however, in contrast to the SRNN, we did not impose neither Dale’s principle nor SFA on the encoder and decoder. Specifically, we assumed the same dynamics and hyperparameters setup for the membrane potential and spike generation.

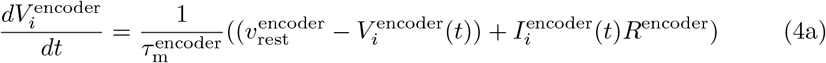

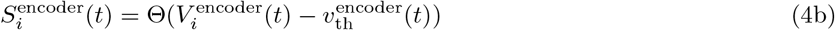

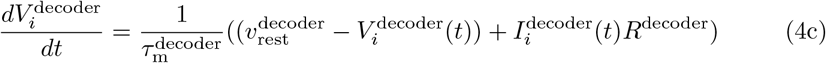

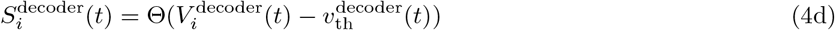

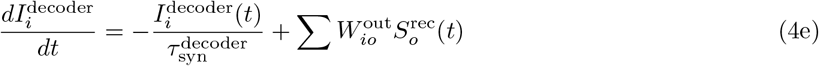

where 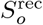 represents the spiking output from the excitatory neuronal population in the SRNN. 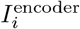 was provided by the continuous-valued cueing or sensory input (Fig. 2B), whereas the dynamics of 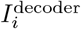 was driven by the output of SRNN weighted by an output matrix ***W***^out^ (Eq. 4e). Finally, the decoder output produced a two-dimensional continuous-valued voltage vector ***V***^decoder^, which represented the probability of making corresponding decisions (e.g., [0, 1] representing the left choice and [1, 0] representing the right choice).

### Imposing biological constraints

#### Dale’s principle

Similar to the previous work [7], we imposed Dale’s principle onto the SRNN. Specifically, cortical neurons have either purely excitatory or inhibitory effects on postsynaptic neurons, and the ratio *ϕ_exc_* of excitatory cortical neurons to all the neurons was 80%.

According to the excitatory (E) and inhibitory (I) populations, the recurrent weight matrix ***W***^rec^ was decomposed into four blocks: {***W***_EE_, ***W***_EI_, ***W***_IE_, ***W***_II_}. The elements in ***W***_EE_ and ***W***_IE_ were all positive, representing the excitatory-to-excitatory and excitatory-to-inhibitory connections, respectively; whereas the elements in ***W***_II_ and ***W***_EI_ were all negative, representing the inhibitory-to-inhibitory and inhibitory-to-excitatory connections, respectively. We initialized 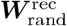 through the Glorot method (Eq. 3), and then rescaled it based on its eigenvalue spectra [1,28]. Specifically, let *ρ* = max {|*λ*_1_|,..., |*λ*_n_|} denote the largest absolute value of eigenvalues of 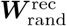; we scaled 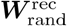 into 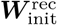 by *g/ρ* such that as

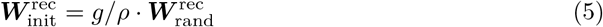

where *g* = 1.5.

We further generated a mask matrix ***D*** = {*D_ij_*} to impose Dale’s principle on 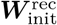 such that 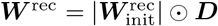, where ⊙ denotes the element-wise product. Specifically, we set *ϕ*_exc_ = 0.8 to reflect the 4:1 ratio of excitatory-to-inhibitory neurons, and assumed that each neuron has its own constraint item

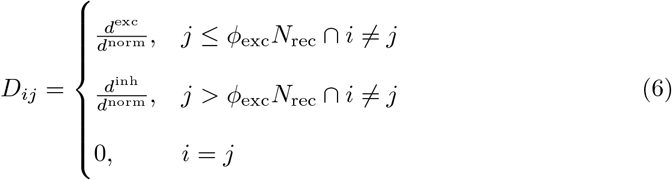

where *d*^exc^ and *d*^inh^ denote the constraints for E and I neurons, respectively; namely, *d*^exc^ was used for ***W***_EE_ and ***W***_IE_, and *d*^inh^ was used for ***W***_II_ and ***W***_EI_; there was no self-connection so that *D_ii_* = 0. Similar to [7, 29], we set *d*^exc^ = 1 and 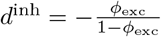 in order to keep the balance of excitation and inhibition (i.e., ∑_exc_ *d*^exc^ + Σ_inh_ *d*^inh^ = 0).

Furthermore, *d*^norm^ is a normalizing constant: 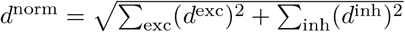.

#### Spike frequency adaptation (SFA)

SFA is a dynamic self-inhibition mechanism that allows neurons adapt their spiking threshold to decrease firing. Recently, it has been shown that SRNNs with SFA can significantly increase the longer short-term memory capability [25]. Briefly, let *A_i_* denote an adaptation term for the spiking threshold. The dynamics of *A_i_*(*t*) was described as follows

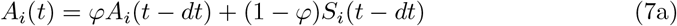

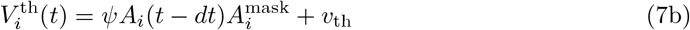

where *ψ* > 0 is a scale parameter. Specifically, when the *i*-th neuron fired, *A_i_* increased instantly; otherwise, it decayed with a factor of 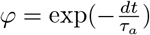 (Eq. 7a), where *τ_a_* = 400 ms denotes the adaptation time constant (*τ*_a_ ≫ *τ*_m_ and *τ*_a_ ≫ *τ*_syn_). Furthermore, the voltage spiking threshold 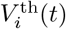 was adjusted based on *A_i_*(*t*) and a binary mask 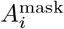. From a total of *N_rec_* neurons, we randomly selected 25% of population and assigned them with SFA 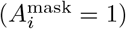. As discussed in [25], neurons with SFA introduce a spread of longer time constants into the SRNN and increase the long short-term memory (LSTM) capacity.

### SRNN training

The decoder output 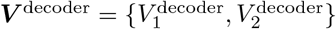 was a two-dimensional continuous-valued vector representing the voltage values from the decoder’s neurons. Let ***y*** = {*y*_1_, *y*_2_} denote the corresponding target vector output; we used the mean-squared error as the cost function 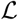, which summed over all single trials loss 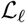 using a batch size *N*_batch_:

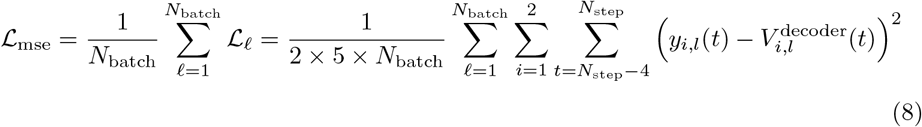

where *ℓ* is the trial index, and *N*_step_ = *T/dt* is the total number of time steps needed within a single trial. We used the last 5 time steps during the sensory presentation period to compute the error.

In addition, we incorporated a firing rate regularization term 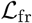 into the cost function, which penalized the mean firing rate of neuronal population. This regularization avoided the excessive firing frequency, also prevented the firing rate of excitatory neurons higher than that of inhibitory neurons. Specifically, the regularized cost function was written as

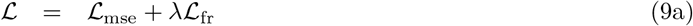

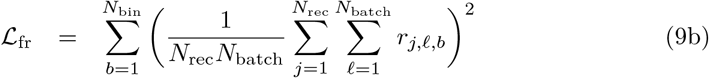

where *r_j,ℓ,b_* denotes the instantaneous firing rate of the *j*-th neuron at the *b*-th temporal bin (1 bin = 5 × *dt* =10 ms and *N_bin_* = *N*_step_/5 = 80) in the *ℓ*-th trial; and *λ* denotes the regularization coefficient.

We further used the Adam algorithm to train all weight matrices of SRNN [30]:

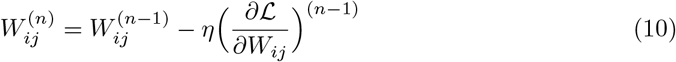

where *η* > 0 is a learning rate parameter, and 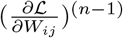 denotes the gradient evaluated on the parameter *W_ij_* at iteration *n* — 1. In BPTT, the computation of gradient back-propagation was defined by the chain rule:

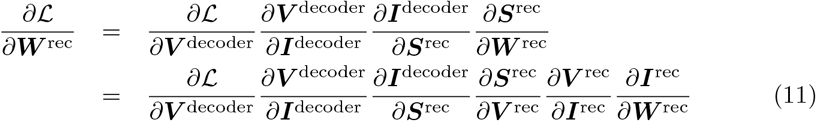

where 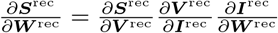 denotes the back-propagated gradient within the SRNN. Therefore, the dynamics of the *i*-th neuron was described by the following equations

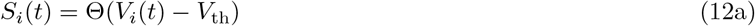

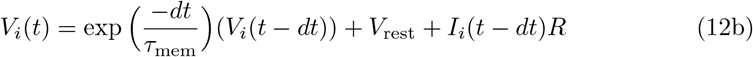

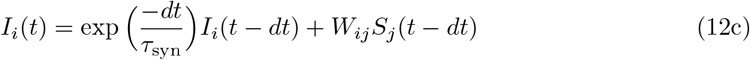

In the chain rule, the corresponding three derivatives were given by

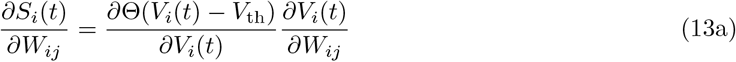

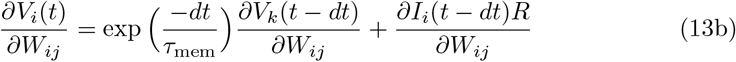

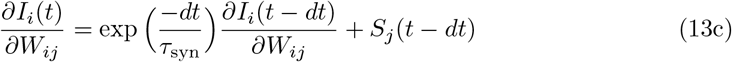

Notably, because of using the Heaviside step function Θ, the gradient information 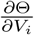 was zero in the absence of spiking [18]. Therefore, the gradient 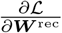 propagated from its previous layers or time steps suffered from the vanishing gradient. To prevent such gradient vanishment, we replaced the gradient in the chain rule by a surrogate gradient known as SuperSpike surrogate gradient [17], which is described in the below.

#### SuperSpike surrogate gradient

The SuperSpike is a nonlinear voltage-based three-factor learning rule, which is capable of training multilayer networks of deterministic LIF neurons to perform nonlinear computations on spatiotemporal spike patterns [17]. Specifically, we computed the SuperSpike surrogate gradient as follows

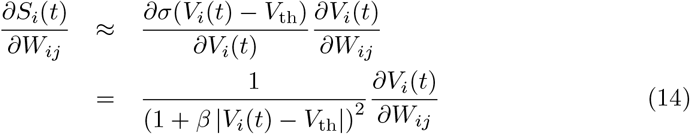

where 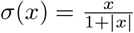 denotes the so-called fast sigmoid function; *β* is a scaling parameter that defines the sharpness of the approximate derivative: the greater *β* is, the sharper the derivative becomes. In Eq. 14, we approximated 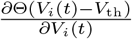 with a smooth surrogate gradient 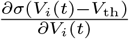.

The computation of SuperSpike gradient has been interpreted as Hebbian coincidence detection [17]. Specifically, the gradient computation of 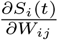 consists of the product of two derivative terms (Eq. 14), the first-order derivative term 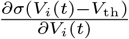 corresponds to the dynamics of the postsynaptic neuron of *W_ij_*. The second partial derivative term 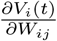 approximates the concentration of neurotransmitters at the synapse *W_ij_*, which represents the presynaptic neuron activity. Thus, training based on SuperSpike gradient is considered as an analogue of Hebbian learning.

### Identification of rule-specific neuronal tuning

Given single-trial spike train simulations of each neuron, we displayed the spike rasters and computed the peri-stimulus time histogram (PSTH) using a 8-ms bin size (i.e., 4 bins). To obtain the smooth tuning profile, the PSTH was convolved with a Gaussian kernel (with kernel width of 4) to create a spike density function based on 50 simulated trials, which was further Z-scored by subtracting the mean firing rate during the delay period (for both rules) and dividing the standard deviation (SD) over the same period. The neurons with Z-scored peak firing rate above a threshold were considered to have sharp tuning. We first screened the neurons with one or more sharp candidate peaks during the delay period for the subsequent analysis. Neurons with very low mean firing rates (< 2 Hz) during the delay period were not considered in the tuning analysis. We used a similar criterion as [19] to identify the genuine tuning peaks. In addition to the minimum firing rate criterion, these candidate peaks needed to have Z-score values of greater than a threshold (e.g., 2.33 for *p* < 0.01). We empirically used a threshold of 2 to match the experimental data. We further categorized the neurons with putative tuning peaks depending on the number of peaks. The majority of identified neurons with rule-specific tuning had 1-2 peaks.

### Characterization of sequential activation

We computed the sequentiality index (SI) to characterize the sequential activation of the population response of excitatory neurons during the delay period. For a given trial, the SI was defined as the sum of the entropy of the peak response time distribution {*P_b_*(*t*_peak_)} (where ∑_*b*_ *P_b_*(*t*_peak_) = 1) of the recurrent neurons and the mean log ridge-to-background ratio of the neurons, where the ridge-to-background ratio for a given neuron was defined as the mean activity of the neuron inside a small window around its peak response time FR_*i*_ (inside peak window) divided by its mean activity outside this window FR_*i*_ (outside peak window) [22, 31]:

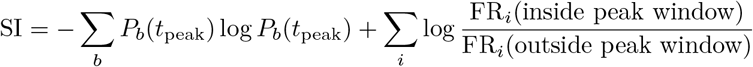

The computer code for computing the SI was available online (https://github.com/eminorhan/recurrent-memory/blob/087a4a67b684ca2e8b6d8102c4221c88105cc16f/utils.py#L16). To test the statistical significance of median SI differences between two conditions, we used a nonparametric rank-sum test throughout the study.

### Dimensionality reduction

To investigate the neuronal population response, we used the principal component analysis (PCA) to extract the principal components of population responses during the delay period. To compute the neural representation or trajectories in the state space, we projected the data onto the subspaces of dominant principal components (PCs). and obtained the low-dimensional latent state trajectory **x**(*t*). Furthermore, we defined the network kinetic energy to characterize the dynamics of SRNN [32]

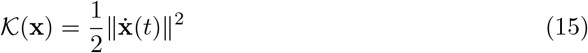

A low value of 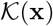 indicates the state reaches a fixed-point region with low or zero momentum.

### Population decoding

We used a long short-term memory (LSTM) recurrent neural network to read out the neuronal population responses during delay period. The goal of the binary LSTM classifier is to predict a category for the temporal sequence. Here we used two LSTM layers for binary classification, and trained the LSTM neural network with BPTT. To avoid overfitting, we used dropout (probability of 0.2) and *L*_2_ regularization (regularization parameter of 0.001). However, the performance of LSTM network was insensitive to the choice of hyperparameters.

Upon completion of SRNN training, we produced a total of 200 trials (evenly for each rule/stimulus configuration) for classification analysis. We temporally binned the spike trains (20 ms bin size) during the delay period. To train the LSTM classifier, we used the PCs as the features. The purpose of PCA was to reduce the number of features and to avoid overfitting. We randomly split 600 trials into two groups, 90% used for training, and 10% used for testing. We conducted 10-fold cross-validation and then computed the average test accuracy. The default classification threshold was 0.5.

### Software implementation

The RNN implementation was built upon the Python package Norse (https://github.com/norse/norse), based on PyTorch and deep learning-compatible spiking neural network components, including network architectures and optimization algorithms. Our custom software implementation can be found in https://github.com/Jakexxh/SNN_PFC-MD.

## Results

### Training SRNN for the 2AFC task

In the default setup (*n* = 500), we used 400 excitatory neurons and 100 inhibitory neurons in the SRNN. We trained the SRNN on the Oracle’s cloud-based GPU-accelerated virtual machine (NVIDIA GPU P100, 16 GB GPU memory, 12 CPU cores, 78 GB CPU memory) within 60 epochs (200 batches per epoch), the performance gradually reached 100% (Fig. S1A,B). Based on different random initial conditions, we repeated the training procedure 10 times and obtained 10 realizations of trained SRNN. Each trained network was treated as one set of independent data. In total, we examined the task representations of 5000 simulated (4000 excitatory and 1000 inhibitory) neurons, and used them in the subsequent analyses. The initial distribution of synaptic weights (absolute value) was close to a uniform distribution. After learning, the weight distribution had a long-tailed behavior, and many of synaptic strengths were close to zeros (Fig. S1C).

### Rule-specific neuronal representations

Upon completion of learning, we first examined the neuronal responses at the single-unit level. We identified task-modulated excitatory neurons that showed sharp peaks during the delay period. We found some excitatory neurons showed rule-specific tunings, with typical differential peaks or peak locations between two rules (Fig. 3A,B). As seen in the spike raster examples, some neurons showed sharp tuning at specific timing for coding one rule, but not the other. The number of peaks was comparable between two rules. A smaller percentage of units were tuned to both rules, with distinct temporal offset in tuning peak (Fig. 3C). This emergent property was reminiscent of the experimental finding in rodent PFC neuronal recordings (Fig. 1B). The ratio of task-modulated or rule-specific tuning excitatory neurons was averaged at 8.1% (over 10 realizations), qualitatively similar to experimental findings [19], and the ratio of excitatory neurons representing rule 1 and rule 2 was roughly equal (4.08% vs. 4.02%). In addition, the averaged firing rates of excitatory and inhibitory neurons during the delay period were 4.15 ± 0.21 Hz and 10.53 ± 0.56 Hz (mean±SEM), respectively. Interestingly, we observed a slightly increase in the firing rate of excitatory neurons during the delay period as compared to the cue period, yet a dramatic firing rate decrease in inhibitory neurons (Fig. 3D).

**Fig 3.**
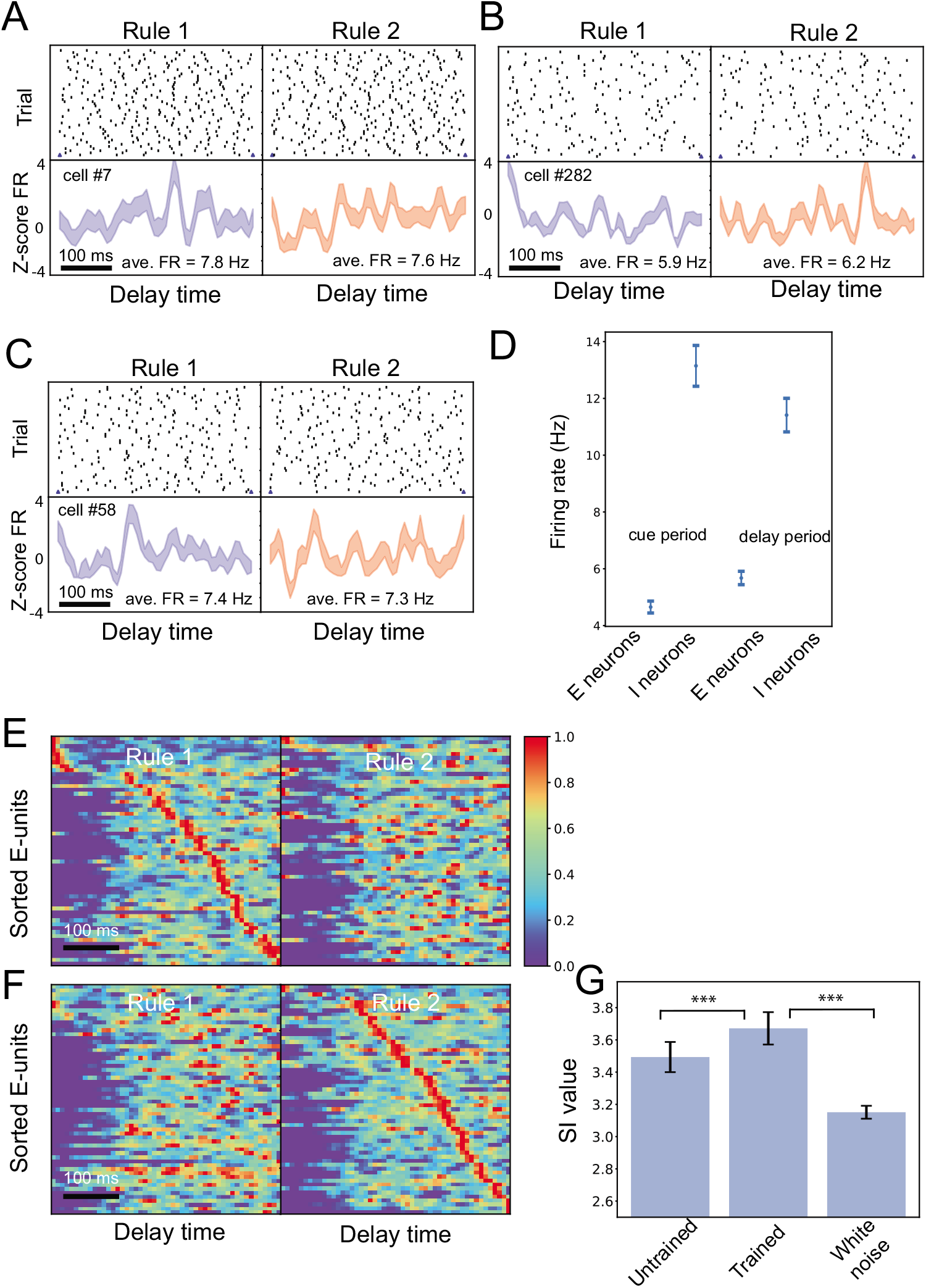
Rule-specific neural representations in trained SRNNs. (A-C) Representative spike rasters and PSTHs of two task-modulated neurons for rule representation. Panel A shows a representative neuron that has sharp tuning to rule 1, but not rule 2; panel B shows a representative neuron that has sharp tuning to rule 2, but not rule 1; panel C shows a representative neuron that has sharp tunings (but at different timings) with respect to two rules. Shaded area denotes SEM. In addition to the Z-score firing rate (FR), the mean FR during delay period is marked in the PSTH panel. (D) Statistics of averaged population firing rates during baseline (fixation) and delay periods. Error bar shows SEM across all neurons. (E,F) Heat maps (each row was normalized between 0 and 1) of normalized task-modulated excitatory neuronal firing rates from one trained SRNN. Note that the sorted E-neurons formed a “neural sequence” for one rule, but not the other. In this example, sequentiality index (SI) was 3.78. (G) Comparison of SI statistics for excitatory neurons between different conditions. Mean±SD statistics were computed from 10 untrained and 10 trained SRNNs. ***, *p* < 10^−3^.

Furthermore, we sorted the rule-specific excitatory neurons according to the peak timing at each rule (Fig. 3E,F). Together, their trial-averaged population responses (visualized as a normalized heat map) showed an emergence of neural sequence (comparing with Fig. 1C). Notably, neurons contributed to different roles in sequence representation at different rules. To quantified the variability of the sequential activation, we further computed the SI measure (Methods) for rule-specific excitatory neurons as well as for the complete excitatory populations (Fig. S2). We also compared the SI statistics between the trained and untrained networks based on the excitatory neuronal population (Fig. 3G). To obtain the chance-level statistic, we fed the white noise input to the trained SRNN and recomputed the SI based on excitatory neuronal activation during the delay the period. The chance-level SI was significantly less than the task condition (*p* = 0.00015, rank-sum test).

In addition, we observed rhythmic firing in spiking activity of a subset of inhibitory neurons during the delay period. This was also independent of their task-tuned properties. A closer look at their PSTHs or auto-correlograms revealed strong beta (15-30 Hz) oscillations at the spiking activity (Fig. 4). In total, 25% (25/1000) of inhibitory neurons exhibited beta rhythms in their simulated spiking activities. Some inhibitory neurons also showed differential rule-specific firing, in terms of oscillatory frequency or phase. However, we did not find any correlation between the power of these rhythmic activity and task performance.

**Fig 4.**
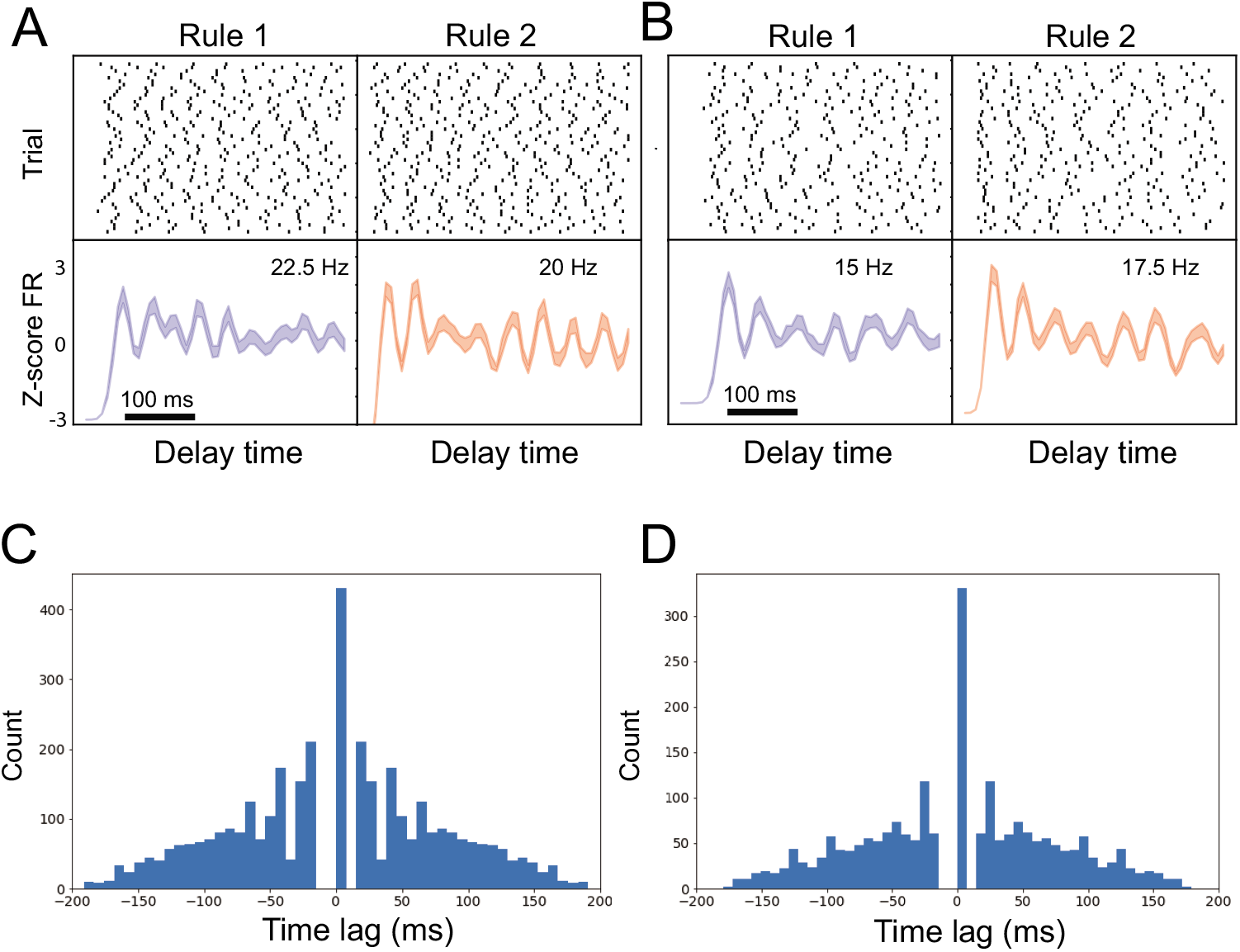
Spiking activities of inhibitory neurons from the trained SRNN showed beta oscillations. (A,B) Two examples of representative spike rasters and PSTHs that showed differential oscillatory frequency or phase in firing between two rules. Error bar represents SEM. The number shows the approximate oscillatory frequency. (C,D) Auto-correlograms of two neurons shown in panels A and B.

### Neuronal coupling and connectivity

Next, we examined the pairwise neuronal coupling, especially those pairs with large (in absolute value) recurrent connection weights of the trained SRNN. In computing the spike-time cross-correlation between E-E, E-I, I-E and I-I neuronal pairs, we focused on the identified rule-tuned neurons as either the trigger or the target. Interestingly, we found many E-E pairs of rule-tuned neurons representing the same rule showed a high peak with a small time lag (2-8 ms) in their cross-correlograms (Fig. 5A). For E-I or I-E pairs, we often found a relatively large trough in their cross-correlograms (Fig. 5B,C). For I-I pairs, the cross-correlogram could have different profiles, depending on whether the rhythmically modulated units were the target (Fig. 5D, left panel), or trigger (Fig. 5D, middle panel), or both (Fig. 5D, right panel).

**Fig 5.**
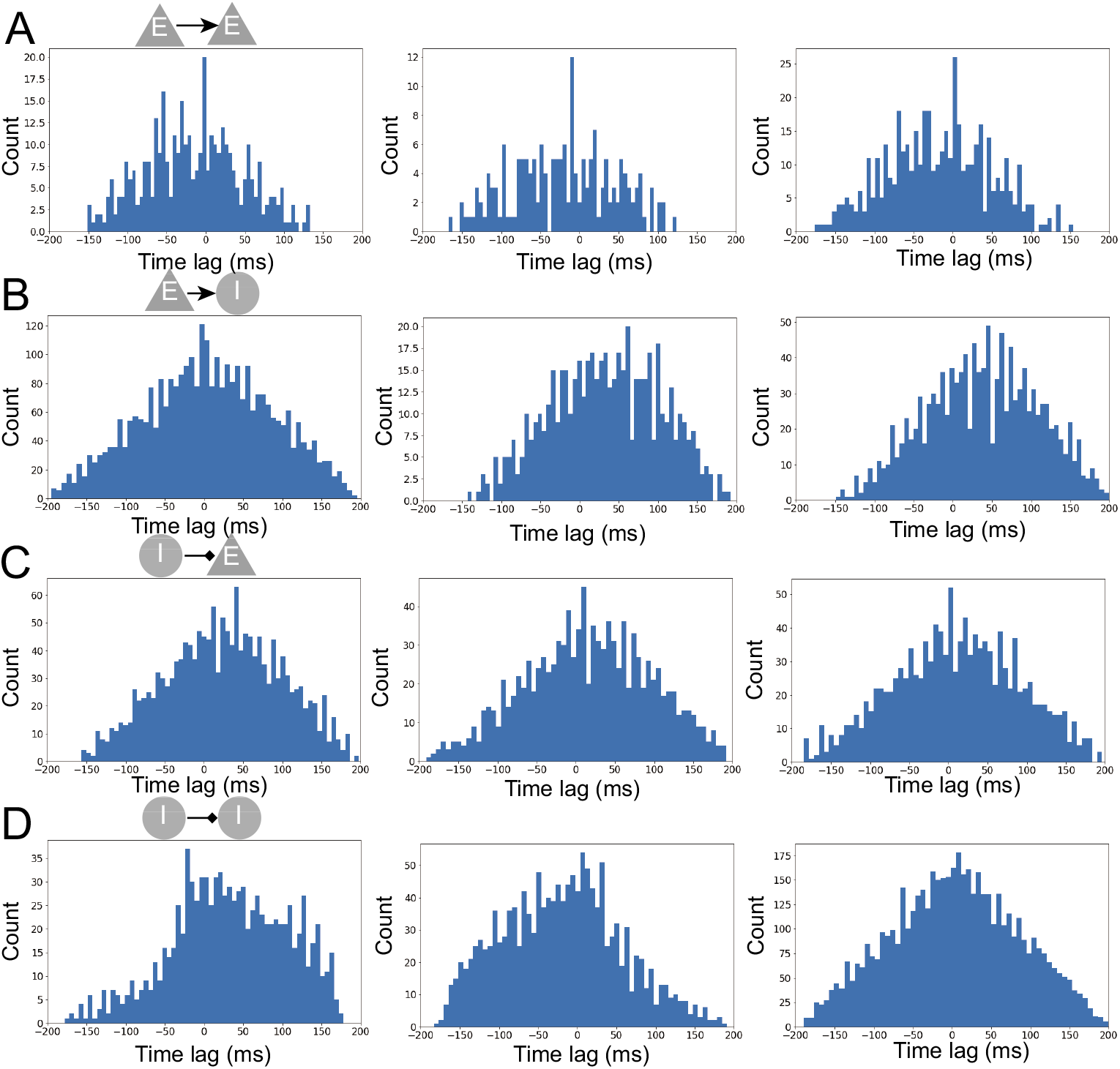
Coupling and cross-correlograms between rule-specific neurons. (A) Three cross-correlogram examples for E-E paired neurons. The trigger and target units are both rule-tuned neurons representing the same rule. (B) Three cross-correlogram examples of E-I paired neurons. The trigger unit is a rule-tuned neuron. (C) Three cross-correlogram examples of I-E paired neurons. The target unit is a rule-tuned neuron. (D) Three cross-correlogram examples of I-I paired neurons.

We further investigated whether single neuronal activities were related to any cluster within the trained network. For each excitatory neuron, we computed the trial variances at three different stages (cue period, delay period, and stimulus presentation period, each period with two rule conditions) and further embedded the 6-dimensional vector in a 2D space using *t*-distributed stochastic neighbor embedding (tSNE) algorithm [33]. Interestingly, we found functionally distinct clusters in the embedded space between rule-tuned and non-tuned excitatory populations (Fig. S3), yet no clear boundary between rule-specific subpopulations.

### Neural trajectory analysis

At the population level, the high-dimensional SRNN dynamics (Fig. 6A) during the delay period can be visualized via dimensionality reduction. Specifically, we conducted PCA and plotted the low-dimensional neural trajectory in the PC space (Fig. 6B). At the PC subspace, we observed the kinetic energy reached the maximum around 100 ms, and then decayed to around zero after 200 ms (Fig. 6C).

**Fig 6.**
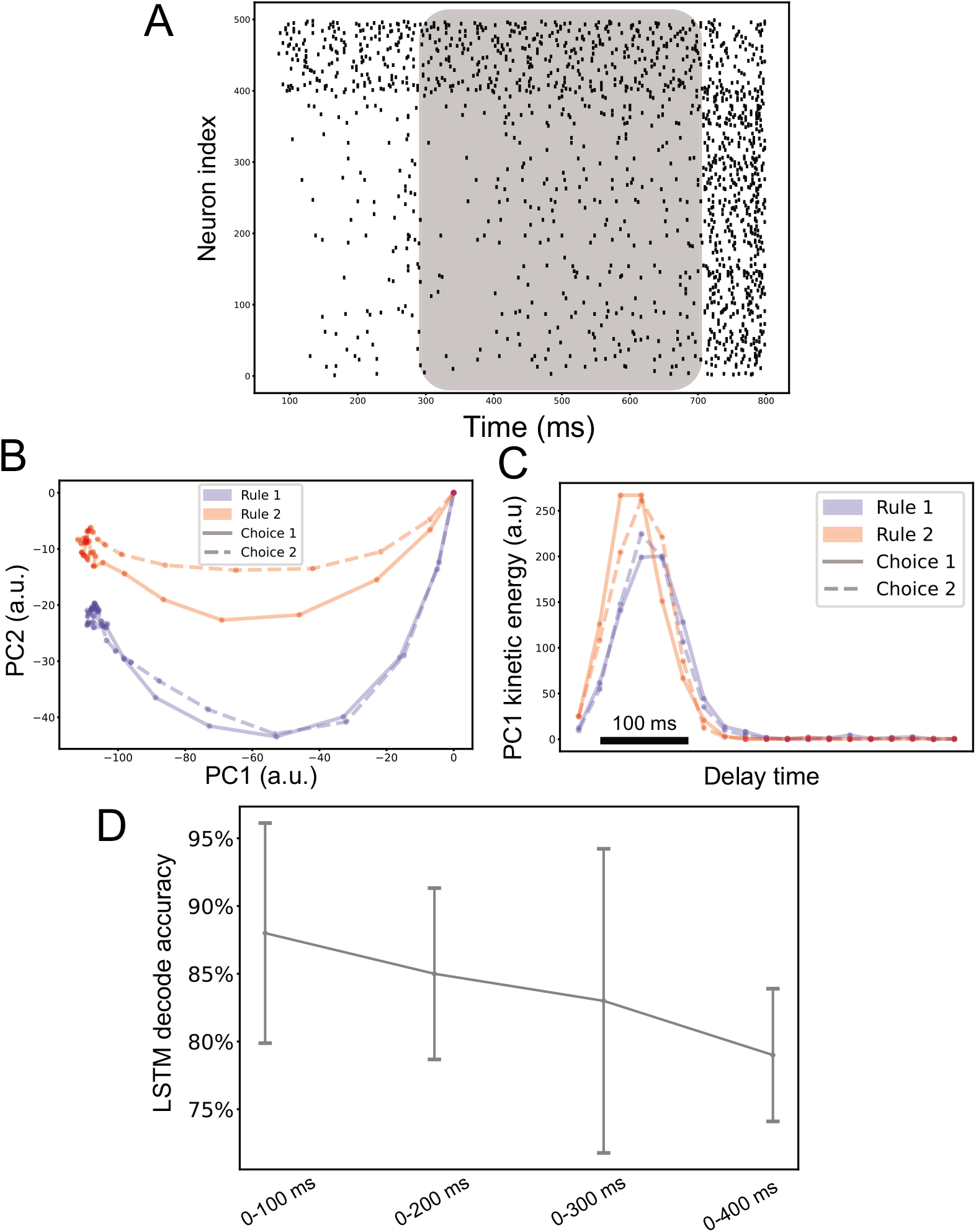
Neural trajectory analysis. (A) Spike rasters of 500 neurons during one single trial (cell #1-400 are excitatory neurons and cell #401-500 are inhibitory neurons). The shaded area denotes the delay period. (B) Neural trajectory ***x*(*t*)** of 400 excitatory neurons projected onto the PC1 and PC2 subspaces (from one trained SRNN) during the delay period. The rule-dependent information (blue vs. red) was maintained and propagated from the start (origin) to the end of delay period. Solid and dashed lines indicate the rule-dependent choices. (C) The curve of kinetic energy 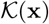 computed from PC1, which shows a high momentum in the first 100 ms, and then gradually decayed to very small value in the last 200 ms. (D) LSTM decoding accuracy (mean±SD) based on 15 dominant PCs of 400 excitatory neuronal firing activity during the delay period.

Next, we ran a population decoding analysis based on 400 excitatory neuronal activities during the delay period. To test the rate code hypothesis, we used an 20-ms non-overlapping moving window to compute the trial-by-trial spike counts of individual neurons. We further projected them onto the PCA subspace, and then decoded the rules from the 15 dominant PCs. We found out that 10-fold cross-validated (on 10% held-out trials) decoding accuracy saturated ~80-90% around 200 ms (Fig. 6D). The performance plateau can be explained by the flat temporal profile of kinetic energy in the last 150-200 ms of delay period. These results also suggest that the additional rule information was more likely to be encoded by individual neurons’ spike timing at a finer timescale beyond the population firing rates.

### Error trials induced by rule encoding uncertainties

Thus far we have limited our analysis on correct trials performed by the trained SRNN. In animal experiments, behavioral errors may arise from executive error, sensory error, or both. The executive error is related to the rule encoding, whereas sensory error is related to the target cue perception. To emulate executive-type error trials, we introduced a mixed representation of two continuous rule inputs by varying their proportional ratios *q* and 1 — *q* (*q* proportion of rule signal 1 plus (1 — *q*) proportion of rule signal 2), and then converted the continuous input via the encoder into a discrete spike train. We fed the spike train that represented the mixed rule signal to the trained SRNN. As expected (Fig. 7A), the network produced an error trial with the highest probability when the ambiguity level was greatest (i.e., *q* = 0.5), and the average network performance gradually decreased when the degree of ambiguity increased.

**Fig 7.**
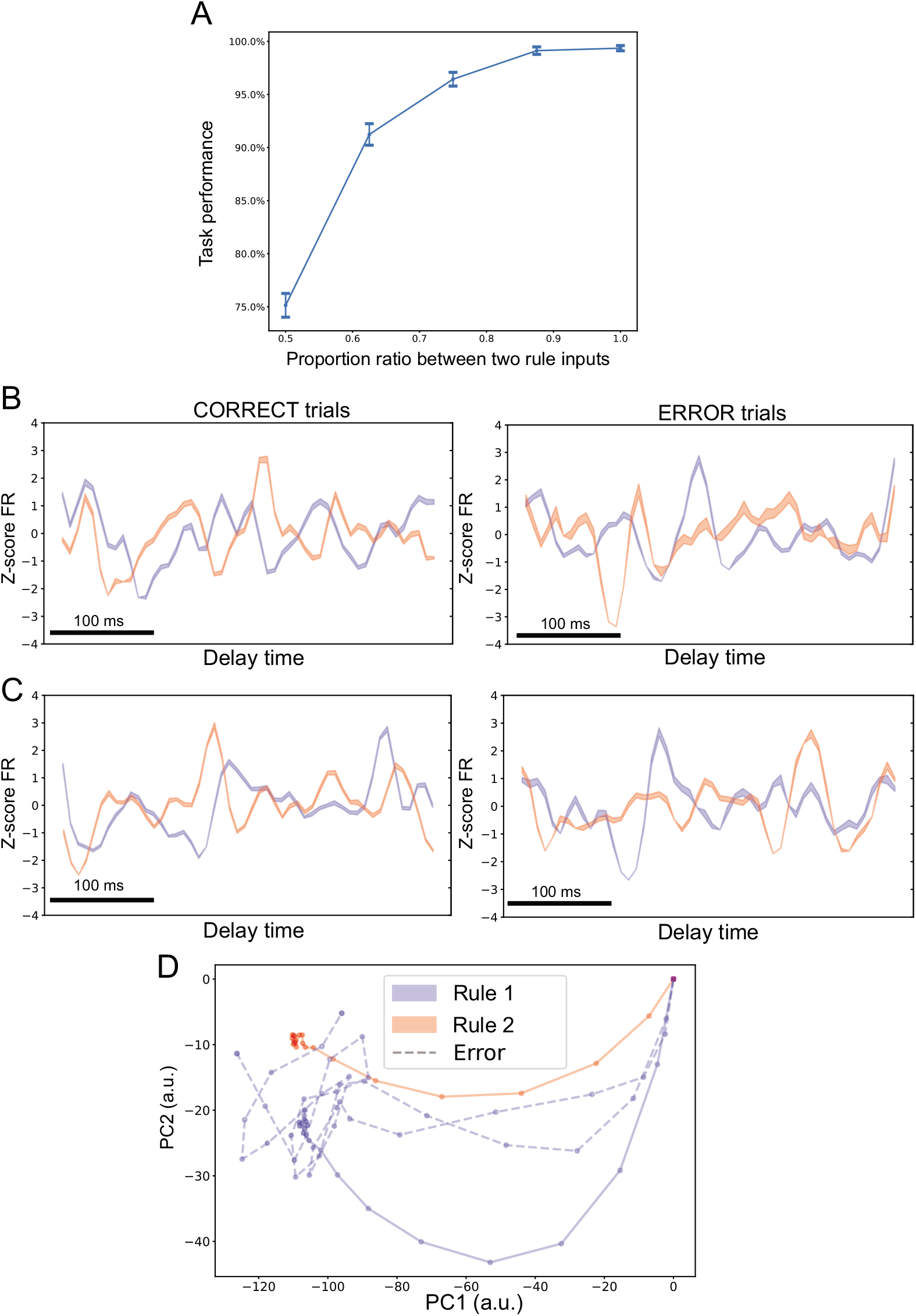
Error trials induced by rule-coding ambiguities. (A) Mean±SD performance curve of SRNN with respect to the mixed rule input with varying proportion ratios. (B,C) Examples of comparative PSTHs of two rule-tuned units between correct and error trials. (D) Projection of two representative error trials (dashed lines, during the delay period) onto the PC subspace yielded neural trajectories deviating from the correct trials. The error-trial trajectories representing rule 1 gradually deviated from the blue trajectory and wandered around at the end of the delay period, creating a higher kinetic energy during the last 200 ms.

We further examined the neural representations of single units during error trials. Generally, we found that the rule-specific tuned units during correct trials changed their firing profiles: (i) units tuned to rule 1 changed preferred tuning to rule 2, or vice versa (e.g., Fig. 7B); (ii) units tuned to both rules changed their peak-firing timing (e.g., Fig. 7C). At the population level, we also projected the neuronal responses of error trials onto the PC subspace; the error-trial trajectory showed a clear deviation from the correct-trial pattern. For instance, the trial mistaken classified as rule 2 started a trajectory closer to rule 1, and then wandered in the neural state space before settling in a region closer to rule 2 (Fig. 7D).

### Impact of elongated delay period

Motivated from the experimental manipulation [19], we extended the 400-ms delay period to 800 ms in the testing phase, and ran the SRNN simulations for the complete 800-ms period. We further examined the neuronal responses in the elongated 400-ms period. Interestingly, we found that some non-task-modulated neurons developed a late peak in the 800-ms delay condition (Fig. 8B); in contrast, the task-modulated neurons that showed an early peak in the first 400-ms delay period preserved the peak timing even in the 800-ms delay (Fig. 8A). The latter case of neuronal responses implies the time-invariant temporal coding. Additionally, we observed a reduction in task performance with an increasing delay period (Fig. 8C), indicating the encoded rule information were gradually lost with increasing time.

**Fig 8.**
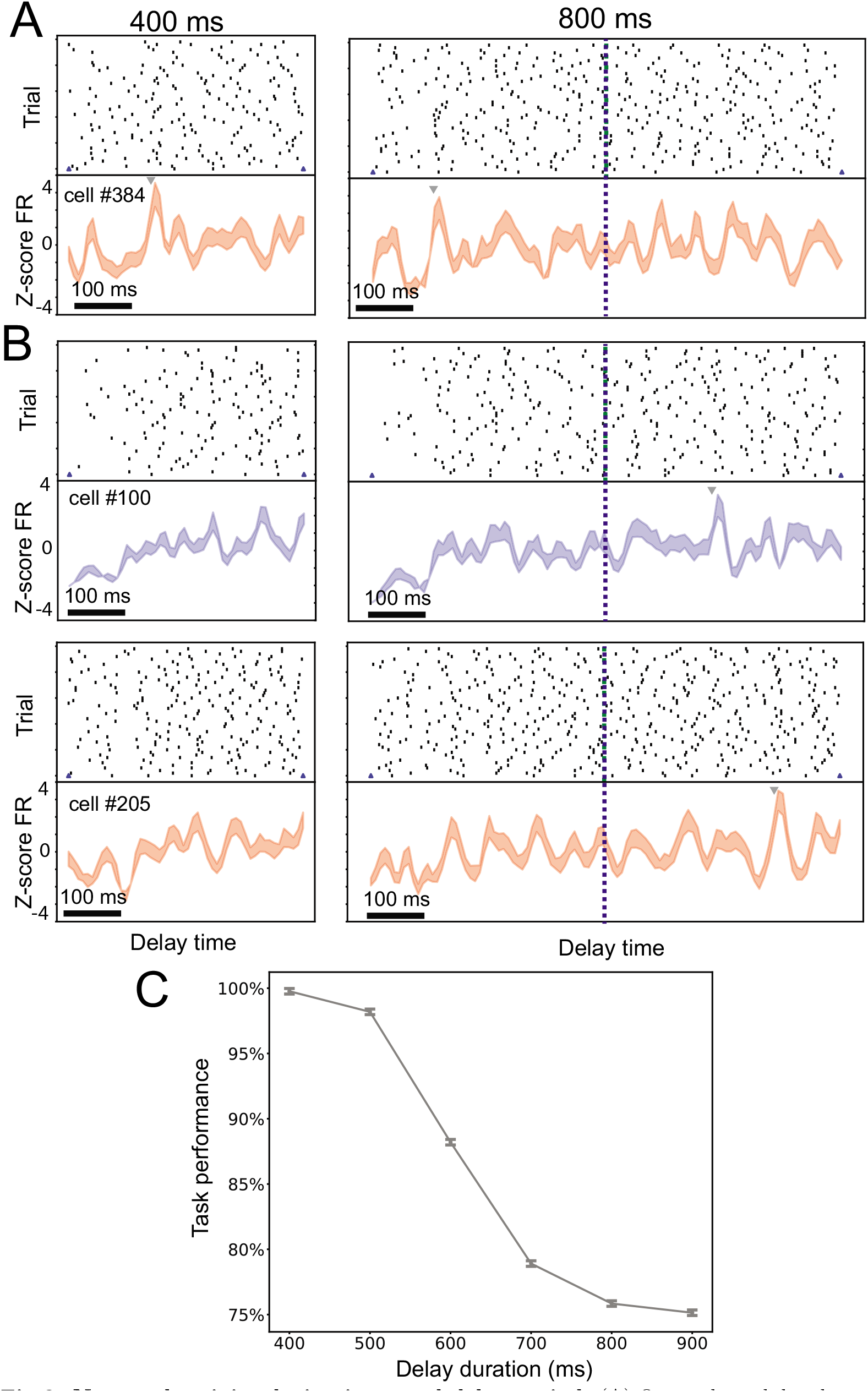
Neuronal activity during increased delay period. (A) One task-modulated neuron preserved the rule-specific peak timing in 800-ms as in 400-ms delay period. (B) New peak timing emerged in the late 800-ms delay period for two non-task-modulated neurons in the original setting. (C) Task performance degraded with an increased delay period.

### Impact of dropped out cortical connections, network connectivity and E/I balance

Furthermore, we investigated the fault tolerance of SRNN with respect to the cortical connectivity. We randomly dropped out a small percentage of synaptic weights in ***W***^rec^ and set them zeros. We ran the network simulations using the modified synaptic weight matrix and tested its impact on neural representations and task performance. We observed the sparsity affected the rule-specific tuning in excitatory neurons. Some observe peak timing disappeared with an increasing sparsity in ***W***^rec^ (Fig. 9A,B). With an increased sparsity level of recurrent weight matrix, we also observed a decrease in task performance (Fig. 9C) as well as cross-validated LSTM decoding accuracy (Fig. 9D). Notably, for the same level of sparsity, the task performance was more robust than the population decoding strategy during the delay period.

**Fig 9.**
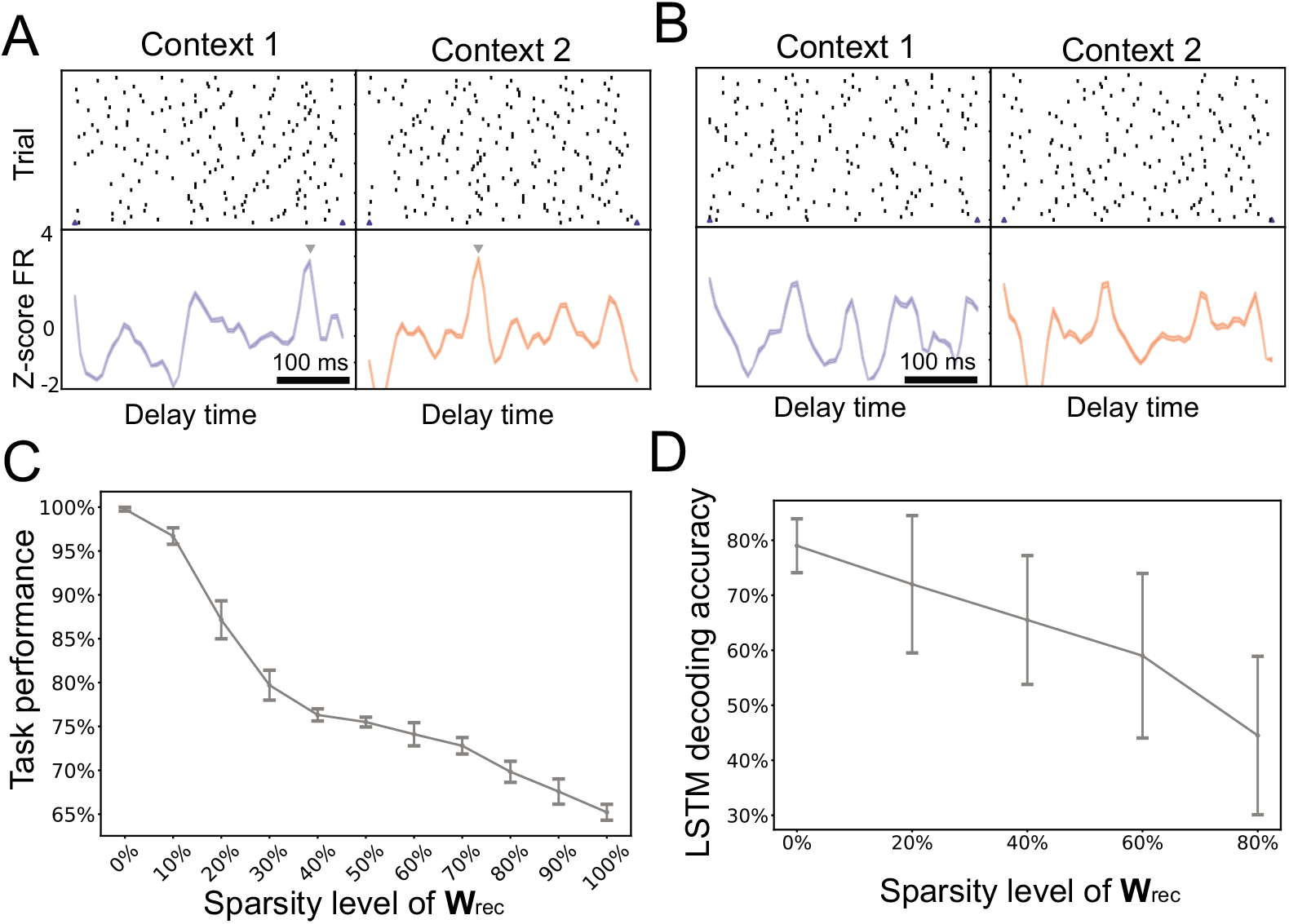
Impact of recurrent weight sparsification on neural representations and task performance. (A) Spike rasters and PSTHs of two task-modulated neurons during the delay period. Arrow marks the peak time. (B) The peak tuning of two neurons in panel A disappeared when ***W***^rec^ was sparsified. (C) Task performance degraded as a result of recurrent weight sparsification. Error bar denotes SD across 10 random realizations. (D) The cross-validated LSTM decoding accuracy degraded with an increasing degree of ***W***^rec^ sparsity. Error bar denotes SD across 10 random realizations.

To examine this effect of cell-type-specific neuronal connectivity on the neural representation and task performance, we scaled up or down the excitatory-to-excitatory ***W***_EE_ connectivity (by multiplying a scaling factor), and measured the changes in firing rates of E and I neuronal populations (Fig. 10A). As expected, the scaling factor affected the E/I firing rate and E/I balance. We further measured the impact on task performance in the trained SRNN (Fig. 10B). When the E/I was imbalanced, the task performance became poor. In the extreme condition where the overall excitation dominated over inhibition, the performance reduced to a chance level. As a comparison, we also scaled up or down the inhibitory-to-excitatory ***W***_EI_ or excitatory-to-inhibitory ***W***_IE_ connectivity and reported the firing rate changes (Fig. 10C,E) and task performance (Fig. 10D,F).

**Fig 10.**
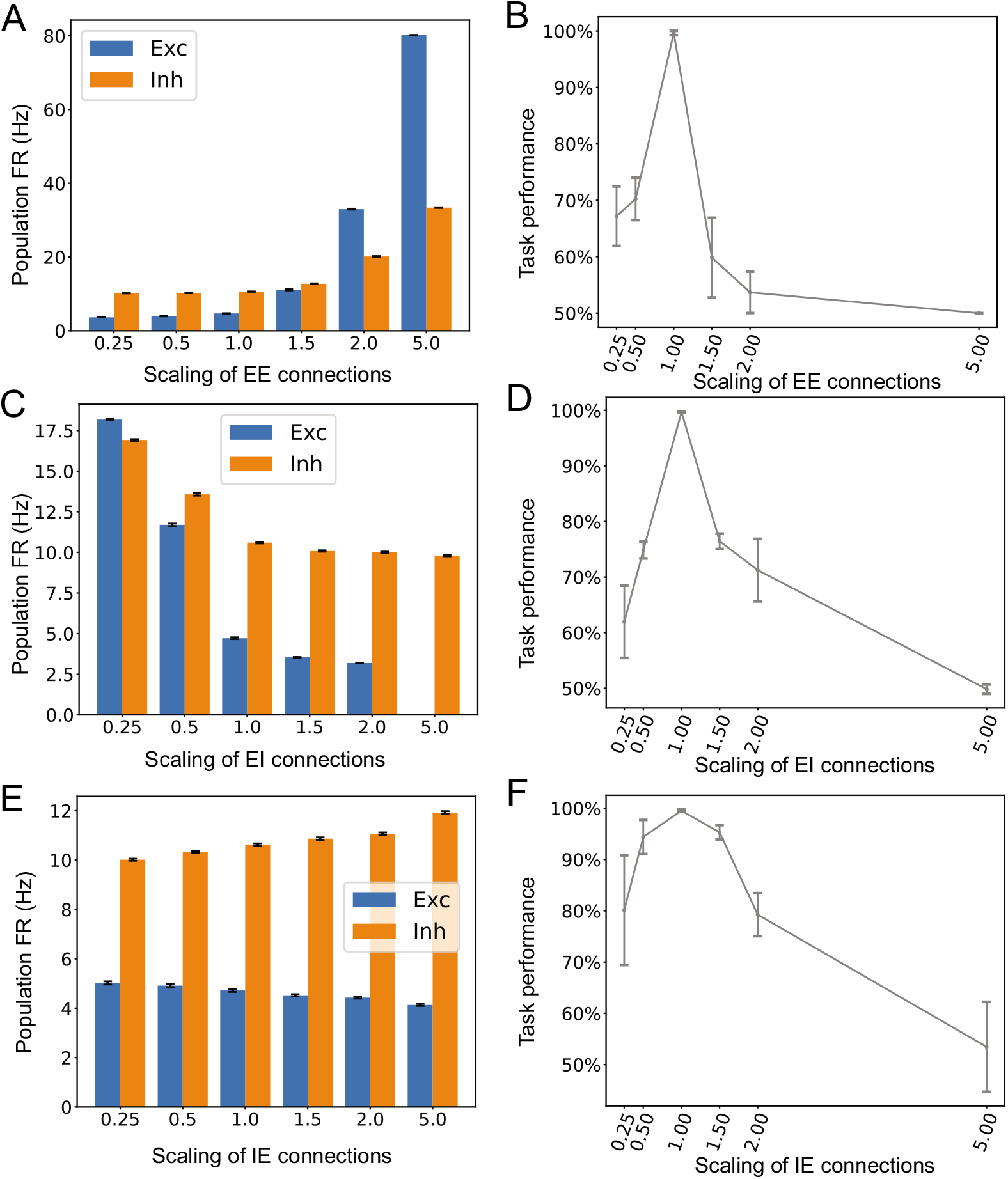
Impact of the cortical connectivity on the task performance and neural representations. (A) Population firing rates (FRs) of excitatory and inhibitory neurons changed with scaled EE connection strengths. Error bar denotes SEM. (B) Scaling up or down ***W***_EE_ changed the E/I level, and further affected the task performance. Error bar denotes the SD over 10 random realizations. (C,D) Similar to panels A and B, except for scaling ***W***_EI_ connection strengths. (E,F) Similar to panels A and B, except for scaling ***W***_IE_ connection strengths.

At the single-unit level, we found that the rule-tuned units decreased their peak firing rates according to the network E/I balance (Fig. S4). Specifically, we observed that the modified excitatory-to-excitatory or inhibitory-to-excitatory connections generated either conflicting or ambiguous rule tuning (Fig. S4A), or diminished rule tuning (Fig. S4B). Notably, our results were similar the reports shown in optogenetic PFC experiments [19].

### Extension and generalization

To investigate the generalization ability of SRNN, we adapted our computer experiments from the 2AFC task to the 4AFC task. Accordingly, we used four neuronal outputs to represent the four possible choices, and the decision was selected based on a softmax function. Since the computer task difficulty was increased, we thereby increased the size of SRNN from *n* = 500 to *n* = 800. We noticed that the the final task performance of the trained SRNN slightly degraded (~95-98%); however, the rule-tuned neural representations still preserved, at both single unit and population levels (Fig. S5).

Additionally, we investigated whether Dale’s principle was required to produce the emerging properties of SRNN. Therefore, we removed the Dale’s principle constraint and retrained the SRNN. Interestingly, we found that most of reported phenomena still held, at both single cell and population levels (Fig. S6). Therefore, this result suggested that the emerging properties (rule-specific tuning and neural sequences) were not dependent on Dale’s principle. On the other hand, we have also implemented a rate-based excitatory-inhibitory RNN model as described in [7] (results not shown). However, we could not reproduce the rule-specific sharp tuning (Fig. 3 and Fig. 4) and cross-correlogram (Fig. 5) at the single-cell level.

## Discussion

The context or rule-dependent task behaviors have been modeled in the literature [2, 7]. Mante *et al.* trained a rate-based RNN model of selection and integration to simulate the the rule-dependent computation in the monkey PFC [2]. The rate-based RNN model qualitatively reproduces the monkey’s behavior as well as the PFC population responses. There are several major distinctions between our model and theirs. First, our model is a SRNN, which uses a different training algorithm. Second, our model imposes biological constraints onto the recurrent weight connections (i.e., Dale’s principle). Song *et al.* extended the rate-based RNN model by imposing similar biological constraints, but their learning framework was fundamentally different from ours [7]. Notably, the distinction of excitatory and inhibition neurons in the SRNN enabled us to examine the impact of cell type-specific connectivity and E/I balance. Third, we introduced an additional biological constraint known as SFA. This also turned out to be important for training SRNN efficiently. Fourth, we imposed a firing rate regularization constraint to encourage the sparsity of population responses.

Our modeling work was strongly motivated based on the previously published data [19]. Our central result is that the trained SRNN performing a 2AFC task can exhibit neural representations and dynamics that resemble the experimental findings of mouse PFC recordings. The resemblance between the SRNN and experimental data was manifested at the levels of single neurons and population responses. Although we didn’t model the neural sequence explicitly, the neural sequence representation emerged from the trained SRNN for the 2AFC task. Although the exact results may depend on the RNN’s hyperparameters (e.g., time constant, noise level, cost function and other simulation parameters), the general phenomena regarding rule-specific tunings and sequence representations are rather robust. Additionally, Dale’s principle is not a necessary condition for those emergent properties. Recent work [31] has suggested that both sequential and persistent activity are part of a spectrum that emerges naturally in the trained rate-based RNN under different conditions; and many factors, such as intrinsic circuit properties, temporal complexity of the task, Hebbian synaptic plasticity, delay duration variability, and structured dynamic inputs, can affect the neural representation of recurrent neurons in the trained network. Our SRNN modeling also confirmed similar findings. We found that the change in sparsity or strength of recurrent weight connections can affect neural representations under various test conditions.

Additionally, our trained SRNN produced emergent beta oscillatory activity, which was in line with the monkey PFC recordings during working memory tasks [34]. According to the dynamic coding hypothesis [35, 36], the oscillatory activity is created by local feedback inhibition shared by local clusters of pyramidal neurons. During attractor activations, the recurrent connections and specific synaptic potentiation produce a slight excitatory bias in the subset of neuronal assemblies. Once they spike, they further activate a new wave of feedback inhibition and turn down the rest of neurons. Overall, our modeling study provides a computational framework to understand a number of PFC findings in cognitive control and working memory. For instance, we observed diverse (dynamic, persistent, and oscillatory) responses of single-unit activity during the delay period. The emerged oscillatory dynamics at the spiking activity seen in the SRNN demonstrate the computer modeling advantages at a fine temporal resolution, which are missing in the rate-based RNN models.

In general, RNN modeling can provide a computational platform to investigate neural representations of cortical circuits in a wide range of cognitive tasks at a fine timescale. For instance, Yang *et al.* trained a rate-based RNN model for performing 20 cognitive tasks that depend on working memory, decision-making tasks, categorization, and inhibitory control [8]. They found that after training, functionally distinct clusters emerged among recurrent neurons that were functionally specialized for different cognitive processes. Learning often gave rise to compositionality of task representation. In addition, the trained network developed mixed task selectivity similar to recorded PFC neurons after learning multiple tasks sequentially. Extension of our SRNN in a continual learning setting [8, 37, 38], will be the subject of future investigation.

The PFC plays a key role in cognitive flexibility, and its neural representation and working memory function are highly dependent on its interaction with the mediodorsal thalamus (MD) [5, 19, 20, 39]. Recently, the MD has been shown to play a modulatory role in cognitive control by augmenting effective connectivity between PFC neurons. In parallel to their experimental circuit dissection, computational models have been developed to account for their interactions [19, 20]; extension of our SRNN framework to modeling biologically-informed PFC-MD network will be the subject of future investigation.

Finally, while our proposed SRNN is biologically plausible, its training procedure relies on supervised learning and back-propagation. However, it remains arguable whether the brain uses back-propagation to perform synaptic modification [40]. In contrast to the error-correcting learning mechanism, Hebbian plasticity and more specifically spike-timing-dependent plasticity (STDP), has become a well-established mechanism for learning. The STDP provides a biologically plausible unsupervised learning mechanism that locally modifies synaptic weights based on the degree of temporal correlations between the pre- and post-synaptic spike events [15, 41-46]. Therefore, applying temporally asymmetric Hebbian learning for SRNN to model neural sequences represents another important future research direction [47, 48]. In our model, the hyperparameters of intrinsic properties (such as the membrane time constant, firing threshold, and reset potential) were fixed. Recently, it has been shown that joint optimization of synaptic weights (i.e., connectivity patterns) and membrane-related parameters (i.e., intrinsic neuronal properties) of SRNN may help perform complex tasks that require information integration and working memory. Future work will further improve the SRNN and validate it in a realistic and complex task setting.

## Acknowledgments

We thank current and former members of the Chen lab for helpful discussion.

## Data Availability

All custom computer software files are available at https://github.com/Jakexxh/SNN_PFC-MD and https://www.cn3lab.org/software.html.

## Author Contributions

Conceived and designed the experiments: ZSC, XX. Supervised the project: ZSC. Provided and collected experimental data: MMH; Performed the computer experiments: XX. Analyzed the data: XX. Contributed the software: XX. Wrote the paper: ZSC, XX.

## Supporting Information

**Fig S1.**
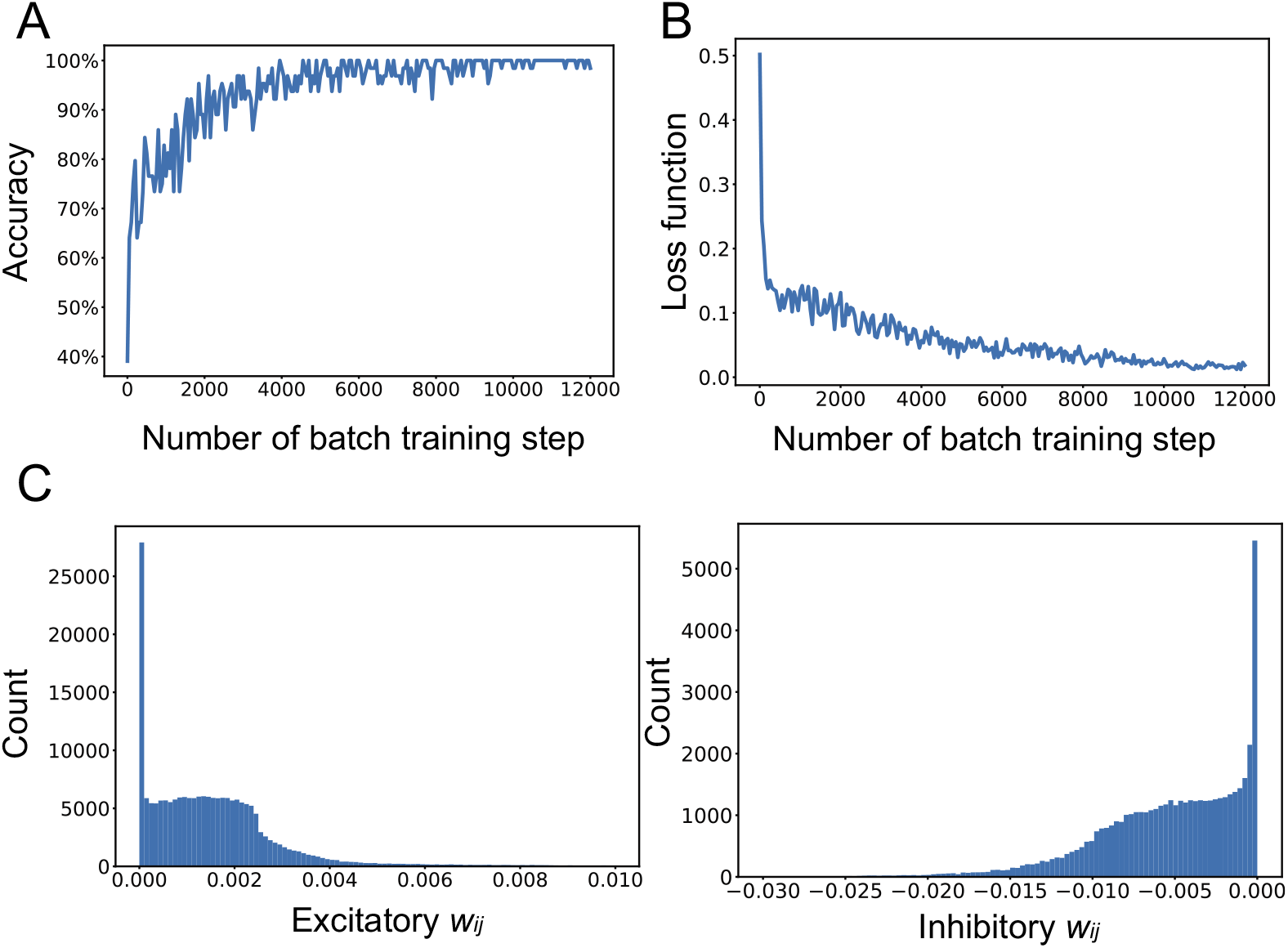
Training SRNN. (**A**) Learning curve for the classification accuracy. (**B**) Learning curve for the regularized loss function. (**C**) Distributions of learned excitatory and inhibitory synaptic connections of a trained SRNN.

**Fig S2.**
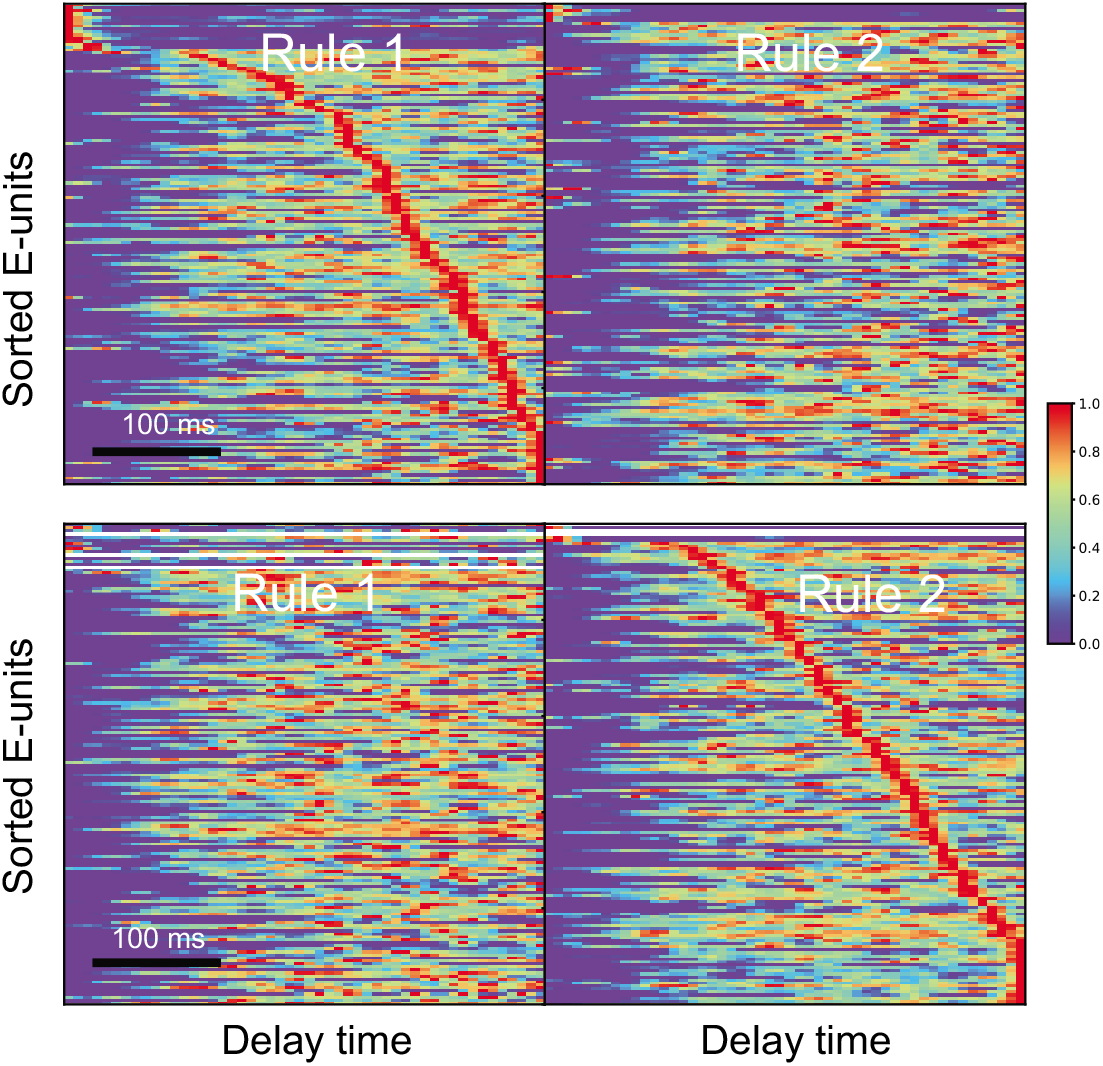
Rule-specific neural sequences. Heat maps of normalized mean firing rates (each row was normalized between 0 and 1) of all excitatory neurons from one trained SRNN.

**Fig S3.**
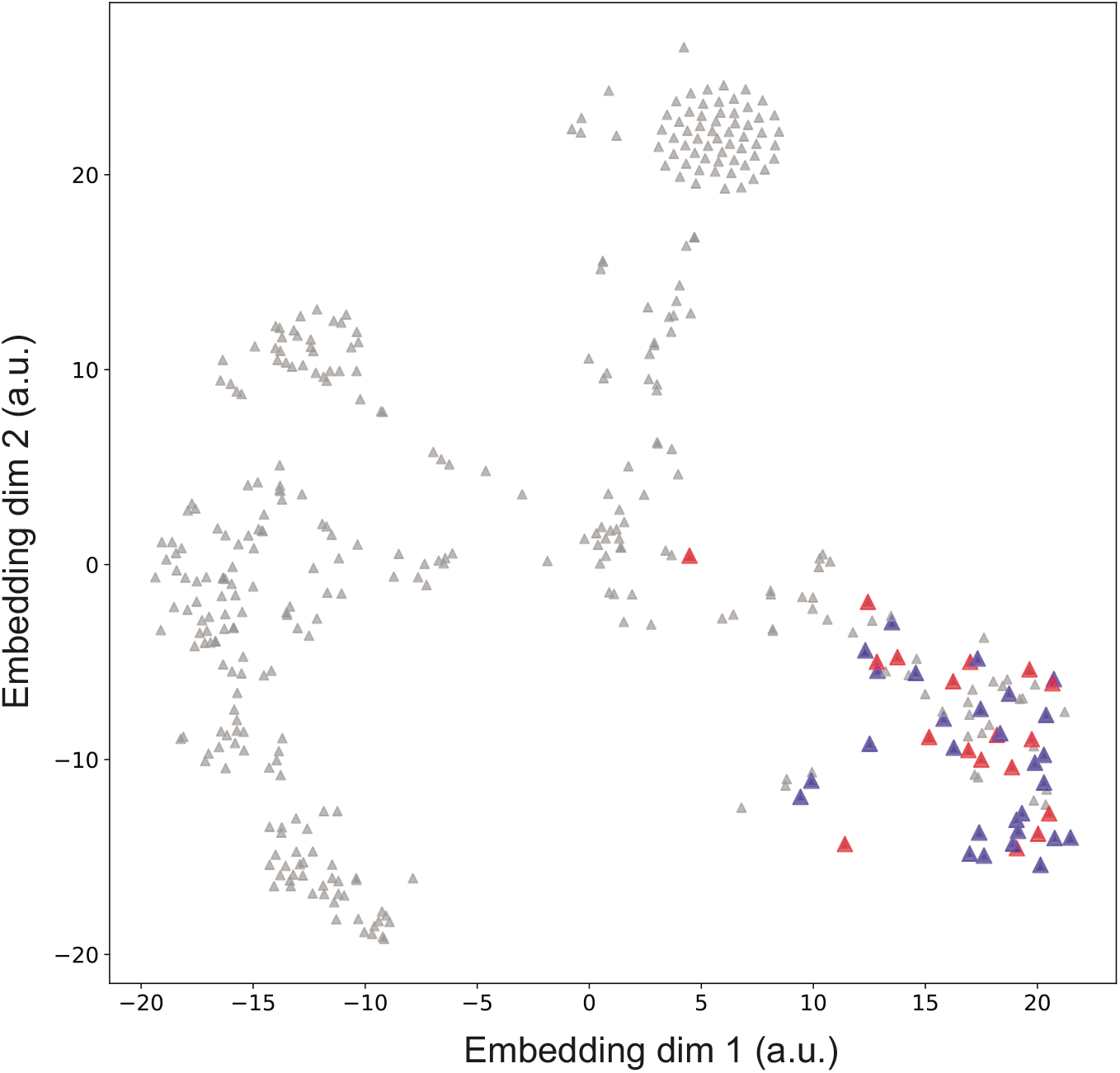
Visualization of functional clusters via stochastic neighbor embedding (tSNE). Each triangle denotes one excitatory neuron, but rule-tuned neurons are shown with bigger markers. Red triangles denote rule 1-specific excitatory neurons, and blue triangles denote rule 2-specific excitatory neurons.

**Fig S4.**
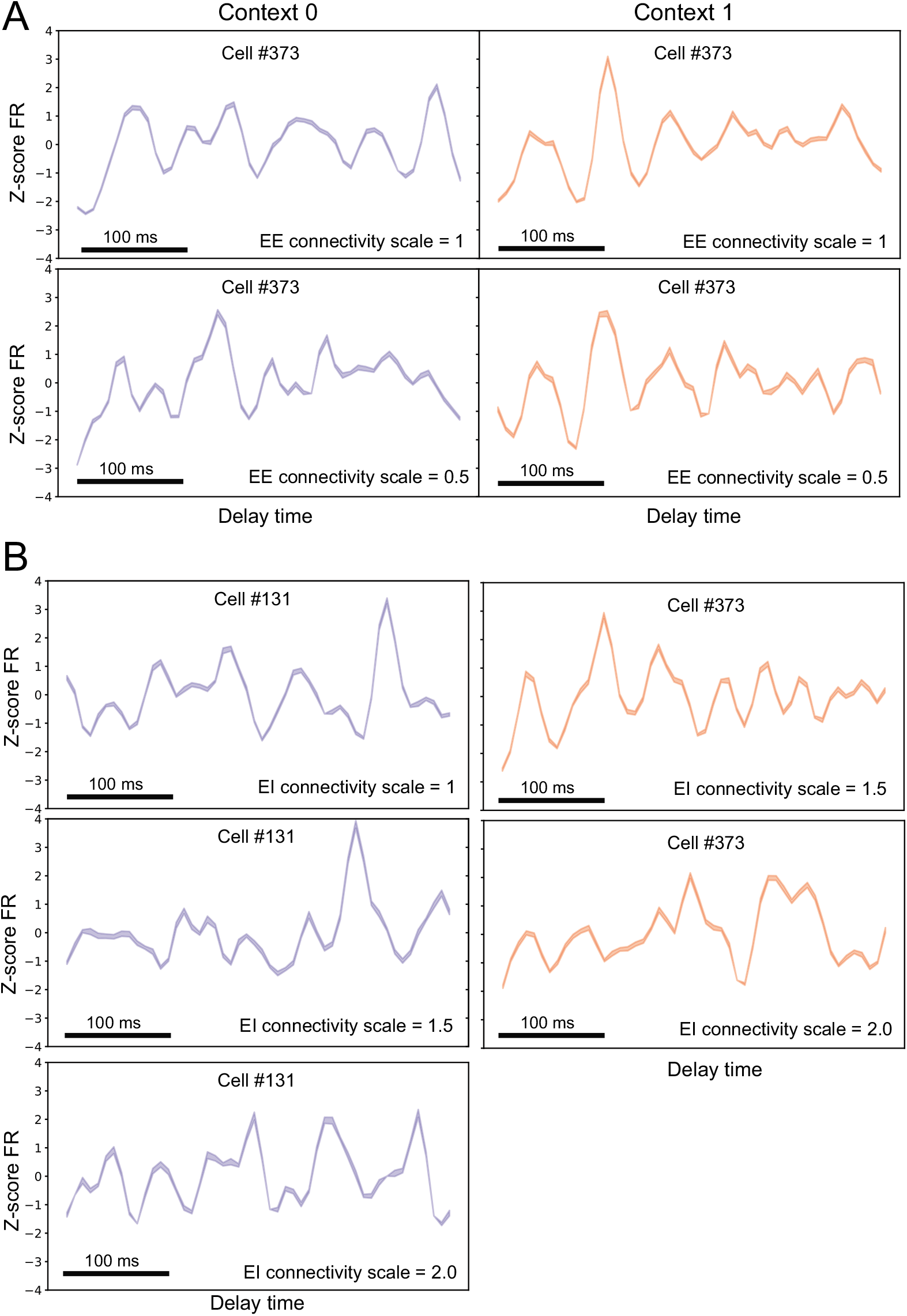
Change in rule-specific tuning due to modified EE or EI connectivity. (A) Change of EE connectivity (scaling of 0.5) generated inappropriate tuning peaks at the opposite rule or reduced the peak tuning. (B) Change of EI connectivity (scaling of 1.5 and 2.0) changed or diminished the peak tuning.

**Fig S5.**
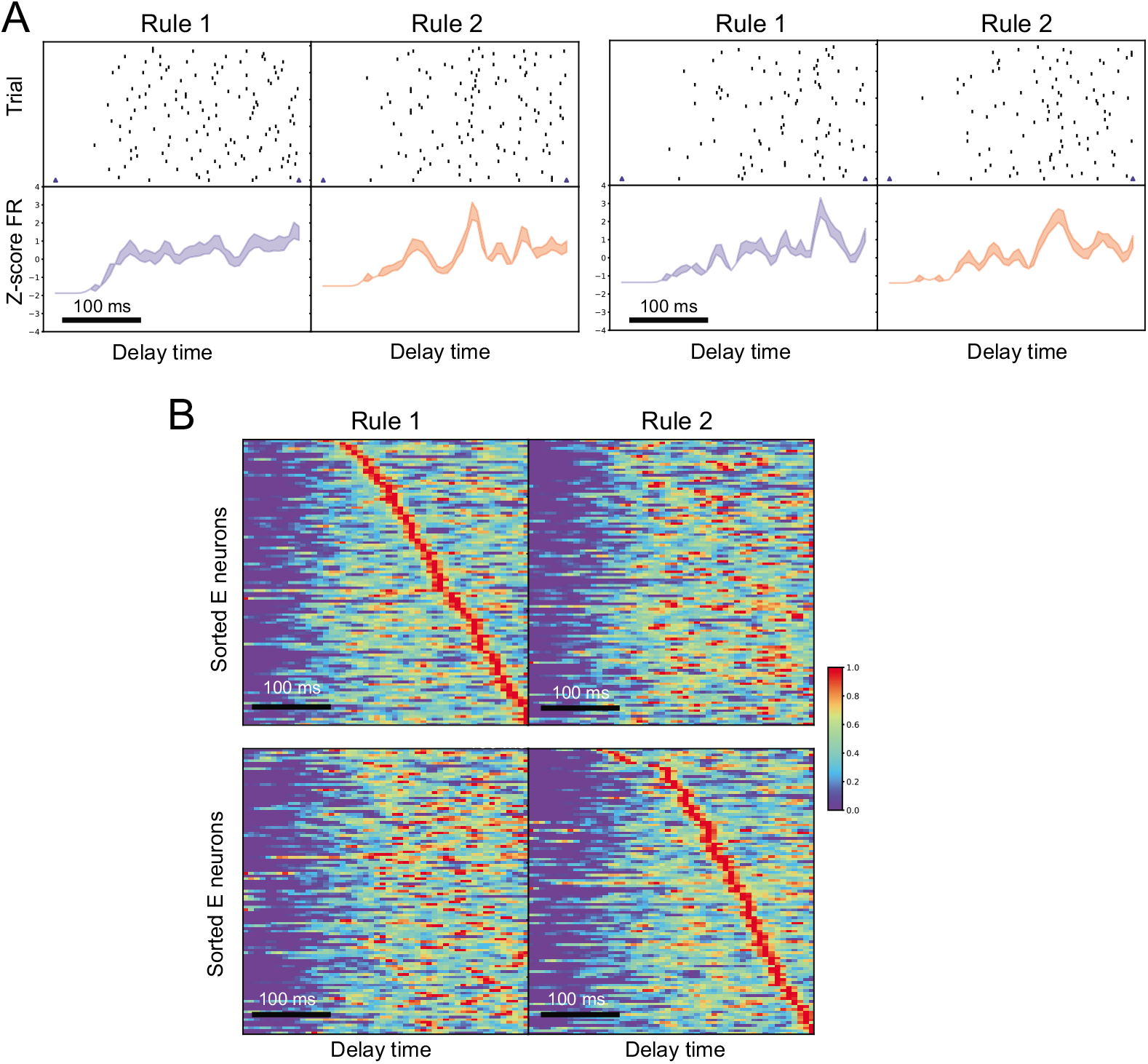
Rule-specific tuning and neural sequence were preserved during the delay period of the 4AFC task. (A) Spike rasters and PSTHs of two representative rule-tuned excitatory neurons. Shaded area denotes SEM. (B) Neural sequence formed by rule-specific excitatory neurons.

**Fig S6.**
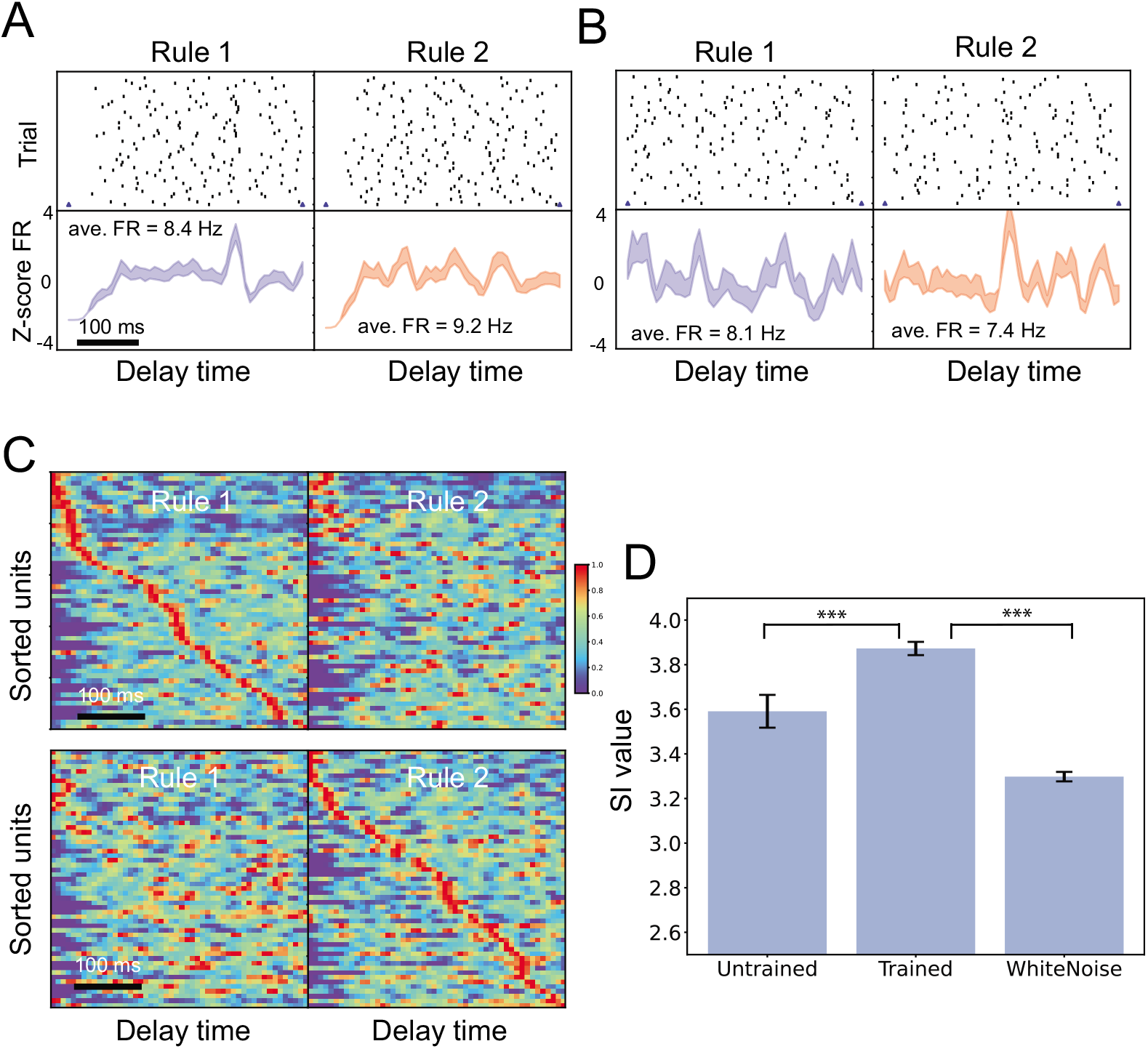
Rule-specific tuning and neural sequence were preserved during the delay period of the 2AFC task, despite the relaxation of Dale’s principle in the SRNN. (A,B) Spike rasters and PSTHs of two representative rule-tuned neurons. Shaded area denotes SEM. Mean firing rate (FR) is marked along the PSTH. (C) Neural sequence formed by rule-specific excitatory neurons. (D) Comparison of SI statistics between different conditions. Mean±SD statistics were computed from 10 untrained and 10 trained SRNNs. ***, *p* < 10^−3^.

## References

1. Sussillo D, Abbott LF. Generating coherent patterns of activity from chaotic neural networks. Neuron. 2009;63(4):544–557.

2. Mante V, Sussillo D, Shenoy K, Newsome WT. Context-dependent computation by recurrent dynamics in prefrontal cortex. Nature. 2013;503:78–84.

3. Sussilo D, Churchland MM, Kaufman MT, Shenoy KV. A neural network that finds a naturalistic solution for the production of muscle activity. Nature Neuroscience. 2015;18(7):1025–1033.

4. Rajan K, Harvey C, Tank D. Recurrent network models of sequence generation and memory. Neuron. 2016;90:128–142.

5. Bolkan SS, Stujenske JM, Parnaudeau S, Spellman TJ, Rauffenbart C, Abbas AI, et al. Thalamic projections sustain prefrontal activity during working memory maintenance. Nature Neuroscience. 2017;20:987–996.

6. Goudar V, Buonomano DV. Encoding sensory and motor patterns as time-invariant trajectories in recurrent neural networks. eLife. 2018;7:e31134.

7. Song HF, Yang GR, Wang XJ. Training excitatory-inhibitory recurrent neural networks for cognitive tasks: a simple and flexible framework. PLoS Computational Biology. 2016;12(2):e1004792.

8. Yang GR, Joglekar MR, Song HF, Newsome WT, Wang XJ. Task representations in neural networks trained to perform many cognitive tasks. Nature Neuroscience. 2019;22:297–306.

9. Zhang X, Liu S, Chen ZS. A geometric framework for understanding dynamic information integration in context-dependent computation. https://www.biorxiv.org/cgi/content/short/20210209430498v1. 2021;.

10. Wolfgang M. Networks of spiking neurons: the third generation of neural network models. Neural Networks. 1997;10(9):1659–1671.

11. Ponulak F, Kasinski A. Supervised learning in spiking neural networks with ReSuMe: sequence learning, classification and spike-shifting. Neural Computation. 2010;22(2):467–510.

12. Sporea I, Grüning A. Supervised learning in multilayer spiking neural networks. Neural Computation. 2013;25(2):473–509.

13. Shrestha SB, Song Q. Adaptive learning rate of SpikeProp based on weight convergence analysis. Neural Networks. 2015;63:185–198.

14. Tavanaei A, Ghodrati M, Kheradpisheh SR, Masquelier T, Maida A. Deep learning in spiking neural networks. Neural Networks. 2019;111:47–63.

15. Panda P, Roy K. Learning to generate sequences with combination of Hebbian and non-Hebbian plasticity in recurrent spiking neural networks. Frontiers in Neuroscience. 2017;11:693.

16. Nicola W, Clopath C. Supervised learning in spiking neural networks with FORCE training. Nature Communications. 2017;8:2208.

17. Zenke F, Ganguli S. Superspike: Supervised learning in multilayer spiking neural networks. Neural Computation. 2018;30(6):1514–1541.

18. Neftci EO, Mostafa H, Zenke F. Surrogate gradient learning in spiking neural networks. IEEE Signal Processing Magazine. 2019;36:61–63.

19. Schmitt LI, Wimmer RD, Nakajima M, Happ M, Mofakham S, Halassa MM. Thalamic amplification of cortical connectivity sustains attentional control. Nature. 2017;545(7653):219–223.

20. Rikhye RV, Gilra A, Halassa MM. Thalamic regulation of switching between cortical representations enables cognitive flexibility. Nature Neuroscience. 2018;21:1753–1763.

21. Fujisawa S, Amarasingham A, Harrison MT, Buzsaki G. Behavior-dependent short-term assembly dynamics in the medial prefrontal cortex. Nature Neuroscience. 2008;11:823–833.

22. Harvey CD, Coen P, Tank DW. Choice-specific sequences in parietal cortex during a virtual-navigation decision task. Nature. 2012;484:62–68.

23. Hardy NF, Buonomano DV. Encoding time in feedforward trajectories of a recurrent neural network model. Neural Computation. 2018;30(2):378–396.

24. Ingrosso A, Abbott LF. Training dynamically balanced excitatory-inhibitory networks. PLoS ONE. 2019;14(8):e0220547.

25. Bellec G, Salaj D, Subramoney A, Legenstein R, Maass W. Long short-term memory and learning-to-learn in networks of spiking neurons. In: Advances in Neural Information Processing Systems (NIPS’18); 2018.

26. Bellec G, Scherr F, Subramoney A, Hajek E, Salaj D, Legenstein R, et al. A solution to the learning dilemma for recurrent networks of spiking neurons. Nature Communications. 2020;11:3625.

27. Glorot X, Bengio Y. Understanding the difficulty of training deep feedforward neural networks. In: Proceedings of the 13th International Conference on Artificial Intelligence and Statistics; 2010. p. 249–256.

28. Rajan K, Abbott LF. Eigenvalue spectra of random matrices for neural networks. Physical Review Letters. 2006;97(18):188104.

29. Okun M, Lampl I. Balance of excitation and inhibition. Scholarpedia. 2009;4(8):7467. doi:10.4249/scholarpedia.7467.

30. Kingma DP, Ba J. Adam: A method for stochastic optimization. arXiv preprint arXiv:14126980. 2014;.

31. Orhan AE, Ma WJ. A diverse range of factors affect the nature of neural representations underlying short-term memory. Nature Neuroscience. 2019;22(2):275–283.

32. Kao JC. Considerations in using recurrent neural networks to probe neural dynamics. Journal of Neurophysiology. 2019;122:2504–2521.

33. van der Maaten LJP, Hinton GE. Visualizing data using t-SNE. Journal of Machine Learning Research. 2008;9:2579–2605.

34. Lundqvist M, Rose J, Herman P, Brincat SL, Buschman TJ, Miller EK. Gamma and beta bursts underlie working memory. Neuron. 2016;90:152–164.

35. Lundqvist M, Herman P, Miller EK. Working memory: delay activity, yes! persistent activity? maybe not. Journal of Neuroscience. 2018;38:7013–7019.

36. Miller EK, Lundqvist M, Bastos AM. Working memory 2.0. Neuron. 2018;100:463–475.

37. Zenke F, Poole B, Ganguli S. Continual learning through synaptic intelligence. In: Proceedings of International Conference on Machine Learning (ICML); 2017. p. 3978–3995.

38. Kirkpatrick J, Pascanu R, Rabinowitz N, Veness J, Desjardins G, Rusu AA, et al. Overcoming catastrophic forgetting in neural networks. Proceedings of the National Academy of Sciences. 2017;114(13):3521–3526. doi:10.1073/pnas.1611835114.

39. Marton T, Seifikar H, Luongo FJ, Lee AT, Sohal VS. Roles of prefrontal cortex and mediodorsal thalamus in take engagement and behavioral flexibility. Journal of Neuroscience. 2018;38:2569–2578.

40. Lillicrap TP, Stantoro A, Marris L, Akerman CJ, Hinton GE. Backpropagation and the brain. Nature Review Neuroscience. 2020;21:335–346.

41. Song S, Abbott LF. Cortical development and remapping through spike timing-dependent plasticity. Neuron. 2001;32:339–350.

42. Fiete IR, Senn W, Wang CZH, Hahnloser RHR. Spike-time-dependent plasticity and heterosynaptic competition organize networks to produce long scale-free sequences of neural activity. Neuron. 2010;65:563–576.

43. Rezende DJ, Gerstner W. Stochastic variational learning in recurrent spiking networks. Frontiers in Computational Neuroscience. 2014;8:38.

44. Lee C, Panda P, Srinivasan G, Roy K. Training deep spiking convolutional neural networks with STDP-based unsupervised pre-training followed by supervised fine-tuning. Frontiers in Neuroscience. 2018;12:435. doi:10.3389/fnins.2018.00435.

45. Lobov SA, Mikhaylov AN, Shamshin M, Makarov VA, Kazantsev VB. Spatial properties of STDP in a self-learning spiking neural network enable controlling a mobile robot. Frontiers in Neuroscience. 2020;14:88. doi:10.3389/fnins.2020.00088.

46. Hao Y, Huang X, Dong M, Xu B. A biologically plausible supervised learning method for spiking neural networks using the symmetric STDP rule. Neural Networks. 2020;121:387–395.

47. Bush D, Philippides A, Husbands P, O’Shea M. Dual coding with STDP in a spiking recurrent neural network model of the hippocampus. PLoS Computational Biology. 2010;6(7):e1000839. doi:doi.org/10.1371/journal.pcbi.1000839.

48. Gillett M, Pereira U, Brunel N. Characteristics of sequential activity in networks with temporally asymmetric Hebbian learning. https://www.biorxiv.org/content/101101/818773v1full. 2019;.

